# Xrn1p Acts at Multiple Steps in the Budding-Yeast RNAi Pathway to Enhance the Efficiency of Silencing

**DOI:** 10.1101/2019.12.12.873604

**Authors:** Matthew A. Getz, David E. Weinberg, Ines A. Drinnenberg, Gerald R. Fink, David P. Bartel

## Abstract

RNA interference (RNAi) is a gene-silencing pathway that can play roles in viral defense, transposon silencing, heterochromatin formation, and post-transcriptional gene silencing. Although absent from *Saccharomyces cerevisiae*, RNAi is present in other budding-yeast species, including *Naumovozyma castellii*, which have an unusual Dicer and a conventional Argonaute that are both required for gene silencing. To identify other factors that act in the budding-yeast pathway, we performed an unbiased genetic selection. This selection identified Xrn1p, the cytoplasmic 5′-to-3′ exoribonuclease, as a cofactor of RNAi in budding yeast. Deletion of *XRN1* impaired gene silencing in *N. castellii*, and this impaired silencing was attributable to multiple functions of Xrn1p, including affecting the composition of siRNA species in the cell, influencing the efficiency of siRNA loading into Argonaute, degradation of cleaved passenger strand, and degradation of sliced target RNA.

## Introduction

RNA interference (RNAi), a gene-silencing pathway triggered by double-stranded RNAs (dsRNAs), can play important roles in viral defense, transposon silencing, and host gene regulation (1,2). In this pathway, dsRNAs originating from viruses, transposons, or cellular loci are initially processed by the ribonuclease (RNase) III enzyme Dicer into ∼21–23-nucleotide (nt) small interfering RNAs (siRNAs) that are paired to each other with 2-nt 3′ overhangs characteristic of RNase III–mediated cleavage. The siRNA duplexes are then loaded into the RNAi effector endonuclease Argonaute, after which one siRNA strand, designated the passenger strand, is cleaved by Argonaute and discarded (or is occasionally discarded without cleavage), thereby generating the mature RNA-induced silencing complex (RISC). Within RISC, the remaining strand, designated the guide strand, pairs with single-stranded RNAs (ssRNAs) to direct the Argonaute-catalyzed slicing of these target transcripts (1,3,4).

Although conserved in most eukaryotes, RNAi is absent from the model budding-yeast species, *S. cerevisiae* (5,6). Indeed, the loss of RNAi has allowed *S. cerevisiae* and some other fungal species the opportunity to harbor Killer, a dsRNA element that encodes a toxin that kills neighboring yeast cells that lack Killer (7). RNAi is present in related budding-yeast species, including *N. castellii* and *Vanderwaltozyma polyspora* (formerly *Saccharomyces castelllii* and *Kluyveromyces polysporus*, respectively), which consequently do not have Killer dsRNA (8,7)*. N. castellii* produces siRNAs that map to hundreds of endogenous loci, a majority of which correspond to transposable elements and Y′ subtelomeric repeats (8). *N. castellii* and other budding yeast thought to have retained RNAi possess a non-canonical Dicer gene (*DCR1*) and a canonical Argonaute gene (*AGO1*), which are, respectively, essential for the production and function of siRNAs in *N. castellii* (8). These two genes are also sufficient for the reconstitution of RNAi in *S. cerevisiae* (8).

In species outside the budding-yeast lineage, factors in addition to Dicer and Argonaute support RNAi pathways. For example, RNA-dependent RNA polymerases (RdRPs) are required for RNAi and related pathways in nematodes, some fungi, and plants (9-15). These enzymes synthesize complementary RNAs either to initiate or to amplify the RNAi response (14,5). Other factors are involved in the loading and maturation of RISC. In Drosophila, R2D2 (named for its two dsRNA-binding domains and association with Dicer-2) associates with Dicer-2, and this heterodimer binds to, recognizes the thermodynamic asymmetry of, and facilitates, with the help of Hsc70/Hsp90 chaperone machinery, the loading of siRNA duplexes into Ago2 (16-22). HSP90 has also been implicated in siRNA loading into AGO1 in plants (23). In human cells TRBP (HIV transactivating response element binding protein), like R2D2 in flies, can help to recruit siRNA-containing Dicer to AGO2 (24,25,19). In the filamentous fungus *Neurospora crassa*, the QIP exonuclease removes the passenger strand during RISC activation (26), and in Drosophila and human, the C3PO endonuclease is reported to have a similar activity that degrades the cleaved passenger strand during RISC maturation (27).

Another factor that can enhance the efficiency of RNAi is the Xrn1 5′-to-3′ exonuclease. Xrn1 is a member of a family of enzymes broadly involved in eukaryotic transcription and RNA metabolism (e.g., processing and degradation) (28). Xrn1 is primarily cytosolic and degrades RNA species possessing a 5′ monophosphate, including mRNA-decay intermediates. Xrn1 orthologs in Arabidopsis and Drosophila enhance RNAi by degrading the 3′ slicing products of RISC (29,30). Likewise, in human cells XRN1 degrades the 3′ slicing fragments and acts together with the exosome to degrade the 5′ slicing fragments of mRNAs targeted by siRNAs (31). By degrading the slicing products of RISC, Xrn1 orthologs likely relieve product inhibition and facilitate multiple turnover of the enzyme. In *Caenorhabditis elegans*, *xrn-1* has also been implicated as being involved in RNAi (32), and its knockdown is associated with the accumulation of certain passenger strands in the microRNA (miRNA) pathway (33), an RNA silencing pathway that derives from the more basal RNAi pathway (34).

Although the only known protein components of the budding-yeast RNAi pathway were Dicer and Argonaute (Dcr1p and Ago1p), we reasoned that, as observed in animals, plants, and other fungi, one or more additional factors might be important for either siRNA production, duplex loading, RISC maturation, or silencing efficiency in yeast. If such a factor did exist, the ability to reconstitute RNAi in *S. cerevisiae* by adding only *DCR1* and *AGO1* indicated that the additional factor either was not essential for RNAi or had other functions in budding yeast, leading to its retention in *S. cerevisiae* after loss of the RNAi pathway. Based on the success of genetic screens and selections in other systems, including Arabidopsis, *C. elegans* and Drosophila, which provided early insight into the core components of the RNAi pathway and identified additional factors that influence RNAi efficacy (35-45), we implemented a genetic selection in *N. castellii* to identify mutants with reduced RNAi activity. This selection identified mutants in the gene encoding Xrn1p. Employing RNA sequencing and biochemical assays, we found Xrn1p acts at multiple steps of the pathway to enhance the efficiency of RNAi in budding yeast. These steps included affecting siRNA populations and loading of siRNA duplexes into Argonaute—steps for which it had not been previously reported to play a role in any species.

## Results

### *XRN1* is a component of the budding-yeast RNAi pathway

A genetic selection was designed to discover novel components of the RNAi pathway in *N. castellii*. This selection centered on an *N. castellii* strain that expressed RNA hairpins that were processed by Dicer to produce siRNAs targeting *HIS3*, *URA3,* and *GFP* mRNAs expressed from exogenous genes that had been integrated into the genome (Figure 1A). The hairpin and *GFP* genes were under the control of the *S. cerevisiae GAL1* and *URA3* promoters, respectively, and the *HIS3* and *URA3* genes, together with their promoters, were obtained from *S. cerevisiae* and replaced their *N. castellii* counterparts in the genome of the selection strain. Because this parental strain was engineered to deploy the RNAi pathway to silence *HIS3* and *URA3*, it did not grow on media lacking histidine and uracil. This strain was mutagenized with ethyl methane sulfonate (EMS), and the cells were plated on media lacking histidine and uracil with the goal of identifying mutants that had a His+ and Ura+ phenotype because they no longer efficiently silenced *HIS3* and *URA3.* As a secondary screen, mutants were examined by flow cytometry for their ability to silence *GFP*, and those that had impaired *GFP* silencing were carried forward for complementation analysis (Figure 1A).

**Figure 1.**
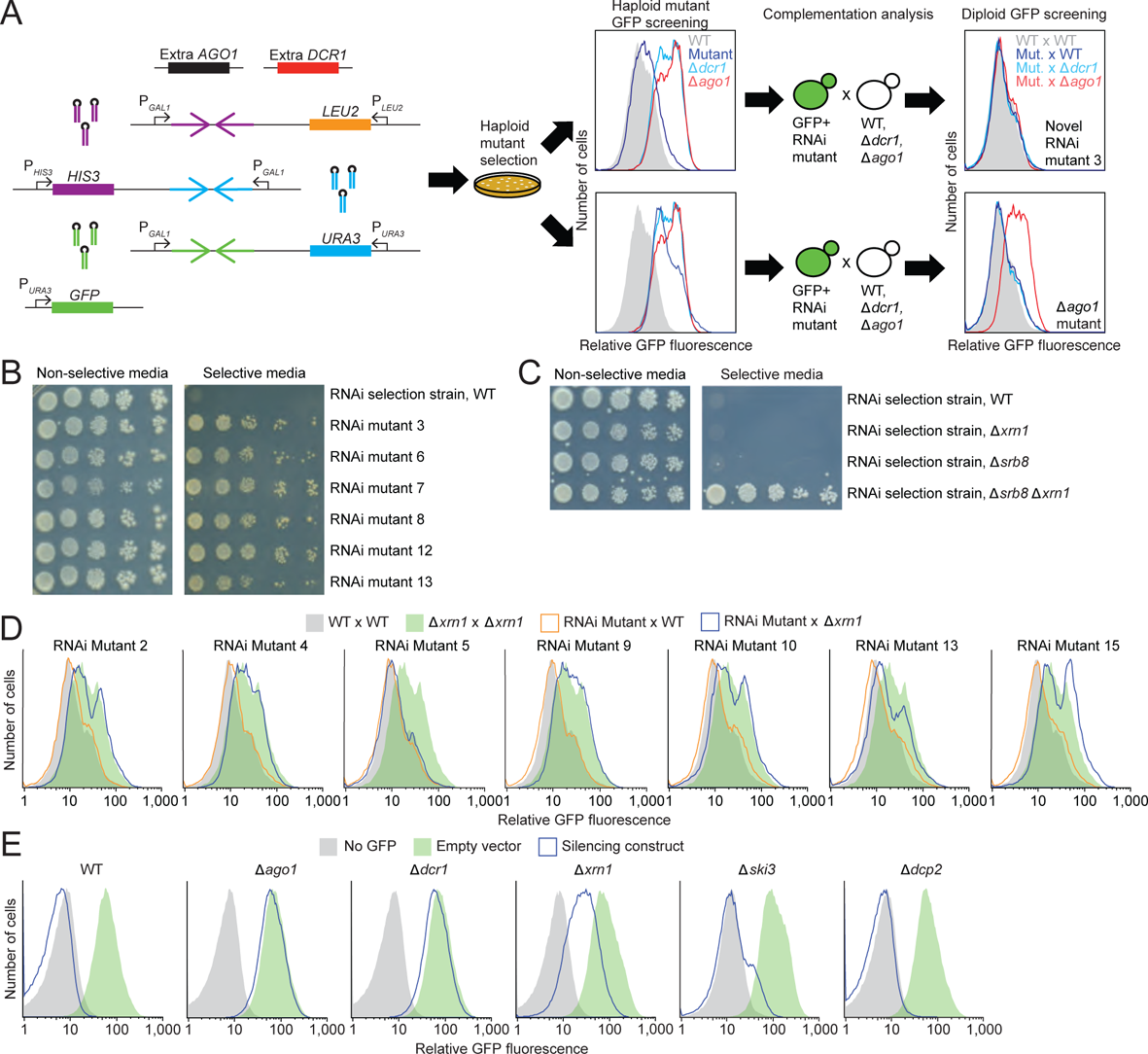
Genetic selection identifying *XRN1* as an enhancer of the RNAi pathway in budding yeast. **A)** Schematic of the genetic selection. At the left are the six genomic insertions of the parental selection strain. Two of these insertions provided an additional copy of the *AGO1* and the *DCR1* genes (black and red boxes, respectively), each added under control of their native promoters. Three of the insertions provided silencing constructs against *HIS3*, *URA3*, and *GFP* genes (purple, blue, and green convergent arrows giving rise to palindromic transcripts that form hairpins) under control of the *S. cerevisiae GAL1* promoter (P_*GAL1*_). These three insertions also provided *S. cerevisiae LEU2*, *HIS3*, and *URA3* genes (orange, purple and blue boxes, respectively) under control of their *S. cerevisiae* promoters, which replaced their endogenous *N. castellii* orthologs. The sixth insertion provided an exogenous *GFP* gene (green box) under control of the *S. cerevisiae URA3* promoter. The haploid selection strain was mutated with EMS, and mutants able to grow on media that lacked Leu, His, and Ura were isolated and screened by flow cytometry for their ability to silence GFP. Mutant strains able to silence GFP were subjected to a complementation analysis in which they were mated with *N. castellii* WT, Δ*ago1*, and Δ*dcr1* strains, and the resulting diploid strains were screened by flow cytometry for their ability to silence *GFP*. Strains for which the ability to silence *GFP* was restored in each of the three matings were carried forward as RNAi mutant strains, with the complementation analysis indicating that they each had one or more recessive mutation that reduced silencing and fell outside of *AGO1* and *DCR1*. **B)** Growth of RNAi mutant strains but not the parental strain on selective media. Representative RNAi mutant strains were serially diluted, plated on either non-selective media (SC (2% galactose)) or selective media (SGal – His, – Leu, – Ura), and allowed to grow for 12 days. For each strain, 1 x 10^5^ cells were plated in the leftmost spot, and dilutions were each ten-fold. **C)** Ability of mutations in both *XRN1* and *SRB8* to confer growth of the selection strain on selective media. Assays compared the growth of single mutant strains (Δ*xrn1* or Δ*srb8*) to that of the double-mutant strain (Δ*srb8* Δ*xrn1*), which were each constructed in the background of the selection strain. Cells were allowed to grow for 10 days; otherwise, as in **B**. **D)** Results of complementation analysis of RNAi mutant strains that had mutations in *XRN1*. Shown are histograms of GFP fluorescence measured by flow cytometry in WT x WT (gray), Δ*xrn1* x Δ*xrn1* (green), RNAi mutant x WT (orange), and RNAi mutant x Δ*xrn1* (blue) diploid strains. All strains were induced. **E)** An intermediate effect on *GFP* silencing observed upon disruption of *XRN1*. Shown are histograms of GFP fluorescence measured by flow cytometry in WT, Δ*ago1*, Δ*dcr1*, Δ*xrn1*, Δ*dcp2* and Δ*ski3* haploid strains that either lacked the *GFP* gene (gray), contained *GFP* but had no silencing construct (green), or contained both *GFP* and a *GFP*-silencing construct (blue). All strains were induced.

Analysis of mutants isolated in pilot experiments illustrated the ability of our selection to find mutants in the RNAi pathway. Mutant strains from these pilot selections were mated to each of three strains: wild-type, Δ*ago1*, and Δ*dcr1*, and the resulting diploids were assessed for their ability to silence *GFP* (Figure 1A). If a new mutation was recessive and in the *AGO1* gene, then the diploid formed with Δ*ago1* would fail to silence GFP, and likewise, if a new mutation was recessive and in the *DCR1* gene, then the diploid formed with Δ*dcr1* would fail to silence GFP. Indeed, these complementation analyses indicated that all of the strains tested had mutations in either *AGO1* or *DCR1*, which showed that the selection scheme worked. To identify additional genes involved in RNAi without having also to contend with a large number of *ago1* and *dcr1* mutants, we modified the selection strain to contain an extra copy of *AGO1* and *DCR1*, each under control of its native promoter, integrated into the genome at an ectopic location.

Mutations that lost silencing were selected in the new parental selection strain (DPB537) with duplicated *AGO1* and *DCR1* genes. As expected, isolated mutant strains but not the parental strain grew on selective media, as assessed by a serial-dilution growth assay (Figure 1B). Each of the isolated strains carried recessive mutations, as assessed from analyses of diploid products of mating with a wild-type strain. Moreover, as illustrated for strain 3 (Figure 1A), 19 strains retained both Ago1p and Dcr1p activities, as indicated by fully restored *GFP* silencing in diploid products of both Δ*ago1* and Δ*dcr1* matings. These 19 strains were carried forward as potentially having mutations in genes encoding another protein needed for efficient RNAi.

To identify mutated genes, we sequenced the genomes of the parental and mutant strains (130-fold and 65–248-fold coverage, respectively), which identified many protein-coding or tRNA changes in each of the mutant genomes (median 47, range 2–120) (Table S1). Of the 19 mutant strains sequenced, seven had a mutation in *XRN1*, the gene for the 5′-to-3′ cytoplasmic exoribonuclease that is highly-conserved in eukaryotes (Table S2). All seven were either G-to-A or C-to-T transition mutations characteristic of EMS mutagenesis, and six of the seven were nonsense mutations. The second most frequently mutated gene was *SRB8*, a component of the mediator complex (46,47), which was mutated in four strains, the mutations in three being transitions and the fourth being a single-base-pair deletion. Two strains had mutations in both *XRN1* and *SRB8*.

To examine the ability of *XRN1* and *SRB8* mutations to confer growth on selective media, we tested the growth of the parental selection strain with single and double knockouts of these genes. The *Δxrn1 Δsrb8* strain grew on selective media, whereas *Δxrn1* or *Δsrb8* mutant strains failed to grow (Figure 1C). The requirement for the *SRB8* mutation can be attributed to the fact that the hairpins used to silence the *HIS3*, *URA3,* and *GFP* genes were each expressed from the *GAL1* promoter. Any mutation that lowered transcription from the *GAL1* promoter would decrease silencing and increase expression of the *HIS3*, *URA3,* and *GFP* genes. A mutation in *SRB8*, as a subunit of the mediator complex, might have provided that reduced hairpin expression. Supporting this idea, *SRB8* is transcriptionally induced in the presence of galactose, as are *SSN2* and *SSN3*, two of the other genes that were disrupted in multiple mutant strains, including strains with *XRN1* mutations (46,48,49). Taken together, these observations suggested that a mutation in *SRB8* (or in either *SSN2* or *SSN3*) created a permissive background, in the context of which a mutation in *XRN1* imparted growth. In this scenario, *XRN1* mutations presumably reduced the efficiency of the RNAi pathway without fully disrupting it.

To test the possibility that loss of *XRN1* function leads to a partial reduction in RNAi efficiency, we determined the ability of *XRN1* mutations to silence *GFP*. Seven haploid *xrn1*-mutant strains from our selection were mated with a haploid Δ*xrn1* strain, and we assessed the ability of the resulting diploid strains to silence *GFP*. The experiment compared the silencing of *XRN1/XRN1*, *XRN1/*Δ*xrn1*, Δ*xrn1*/Δ*xrn1* with Δ*xrn1*/*xrn1*-1–7. Six of these seven diploid strains had impaired *GFP* silencing (Figure 1D). These six each derived from a mutant strain with a nonsense mutation in *XRN1*, whereas the one with efficient *GFP* silencing derived from the strain with a missense mutation in *XRN1* (RNAi mutant 5). These results indicated that the missense mutation probably did not impair Xrn1p activity but that the *XRN1* nonsense mutations were each loss-of-function mutations and were at least partially responsible for the reduced *GFP* silencing observed in the haploid mutant strains. Indeed, the Δ*xrn1*/Δ*xrn1* diploid that we constructed, which had no other confounding mutations, also had reduced silencing.

We also examined the consequence of inactivating *XRN1* in an RNAi-reporter strain containing an integrated *GFP* gene and an integrated gene expressing stem-loop transcripts that can be processed into siRNAs that target *GFP*. Compared to deleting *AGO1* or *DCR1*, deleting *XRN1* produced an intermediate *GFP*-silencing defect (Figure 1E), which supported the conclusion that Xrn1p functions in the budding-yeast RNAi pathway but, unlike Dcr1p and Ago1p, is not an essential component of the pathway. In contrast, disrupting other genes involved in cytoplasmic mRNA decay, including *DCP2* or *SKI3*, which code for proteins important for decapping and 3′-to-5′ degradation, respectively (50,51), had minimal effect on silencing *GFP* (Figure 1E), indicating that their respective decay activities are not required for efficient RNAi in budding yeast.

### Xrn1p and Ago1p physically interact and colocalize

As an orthogonal approach to identify components of the budding-yeast RNAi pathway, we expressed epitope-tagged Ago1p in *N. castellii* and used mass spectrometry to identify co-immunoprecipitating proteins. Xrn1p was the most significantly enriched protein in both biological replicates of this experiment (Table S3), suggesting that Xrn1p physically interacts with Ago1p in these cells. This physical interaction was supported by reciprocal co-immunoprecipitation of Ago1p and Xrn1p as assayed by immunoblotting (Figure 2A). Co-immunoprecipitation was at least partially retained in the presence of RNase, which indicated a protein–protein interaction (Figure 2A).

**Figure 2.**
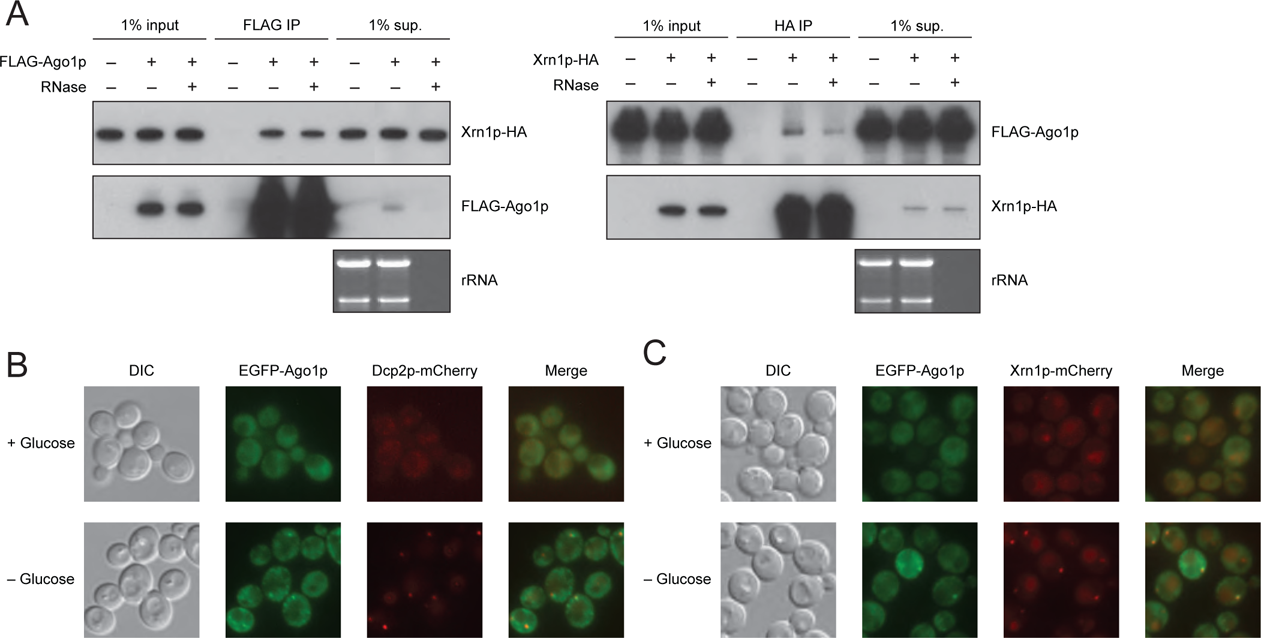
Physical interaction and colocalization of Xrn1p and Ago1p in *N. castellii*. **A)** Physical interaction of Xrn1p and Ago1p in *N. castellii*. Immunoblots show the results of co-immunoprecipitations of FLAG-tagged Ago1p (FLAG-Ago1p, left) and HA-tagged Xrn1p (Xrn1p-HA, right). The samples on the blots were from 1% of the input into the immunoprecipitation (1% input), the immunoprecipitation with anti-FLAG or anti-HA antibodies (FLAG-IP and HA-IP, respectively), and 1% of the remaining supernatant after immunoprecipitation (1% sup.). The immunoprecipitations were performed with and without treatment with an RNase A/T1 cocktail. **B)** Co-localization of Ago1p and Dcp2p observed in *N. castellii* grown in the presence and absence of glucose. Fluorescence microscopy images show the location of EGFP-tagged Ago1p and mCherry-tagged Dcp2p. **C)** Co-localization of Ago1p and Xrn1p observed in *N. castellii* grown in the presence and absence of glucose. Fluorescence microscopy images show the location of EGFP-tagged Ago1p and mCherry-tagged Xrn1p.

To examine the localization of Ago1p and Xrn1p in *N. castellii* cells, we tagged the proteins with GFP and mCherry, respectively. In *S. cerevisiae* grown under standard conditions Xrn1p is reported to be diffusely cytoplasmic, but upon glucose starvation it preferentially localizes to P-bodies, along with other known RNA-decay components including Dcp2p (52). In *N. castellii*, we observed similar behavior of diffuse localization and P-body localization in normal conditions and glucose starvation, respectively, for not only Xrn1p but also Ago1p (Figure 2B and 2C). The colocalization of Ago1p and Xrn1p in *N. castellii* further supported the conclusion that Xrn1p plays a role in RNAi.

### *XRN1* affects small RNA populations in budding yeast

To learn whether Xrn1p impacts the levels of siRNAs in the cell, we sequenced small RNAs from WT and Δ*xrn1 N. castellii* strains after adding known quantities of synthetic standards to each sample, which enabled quantitative comparisons between samples. Mapping of the 22–23-nt reads to previously identified siRNA-producing loci (8) and normalizing to the synthetic standards showed that siRNA levels from the most prolific loci were reduced upon *XRN1* disruption (Figure 3A). Many of these prolific loci were inverted repeats (i.e., palindromic loci), which produce transcripts with a hairpin (stem-loop) structure. For nearly all of the palindromic loci, fewer siRNAs accumulated in the Δ*xrn1* strain (median change, 3.5-fold) (Figure 3A). In contrast, siRNAs mapping to Y′ subtelomeric elements and other non-palindromic loci, in which convergent overlapping transcription produces transcripts that base pair to form bimolecular siRNA precursor duplexes, typically increased in the Δ*xrn1* strain (median change, 4.1-fold). For example, siRNAs from the Y′ subtelomeric elements increased by over 5-fold in the *XRN1*-disrupted strain (Figure 3A). Subtelomeric siRNAs also accumulate in *S. pombe* upon the deletion of its *XRN1* ortholog, *exo2* (53). When classifying the 22–23-nt reads as originating from either palindromic or non-palindromic loci, the proportion of siRNA reads originating from palindromic loci decreased by half in the *XRN1*-disrupted strain (from 74.1% to 37.0%) and the siRNA reads that mapped to non-palindromic loci increased (from 21.5% to 60.8%) (Figure 3B). The disparate changes for the different types of siRNAs observed by sequencing were also observed by probing an RNA blot analyzing small RNAs from the same strains (Figure 3C). This blot showed that an siRNA deriving from a palindromic region decreased by > 2-fold in the Δ*xrn1* strain, whereas a Y′ subtelomeric siRNA increased in expression by > 4-fold. The Y′ subtelomeric siRNA also increased in Δ*dcp2* and Δ*ski3* strains but not to the same degree as in the Δ*xrn1* strain, whereas the palindromic siRNA increased in the Δ*ski3* strain and did not substantially change in the Δ*dcp2* strain (Supplemental Figure 1A).

**Figure 3.**
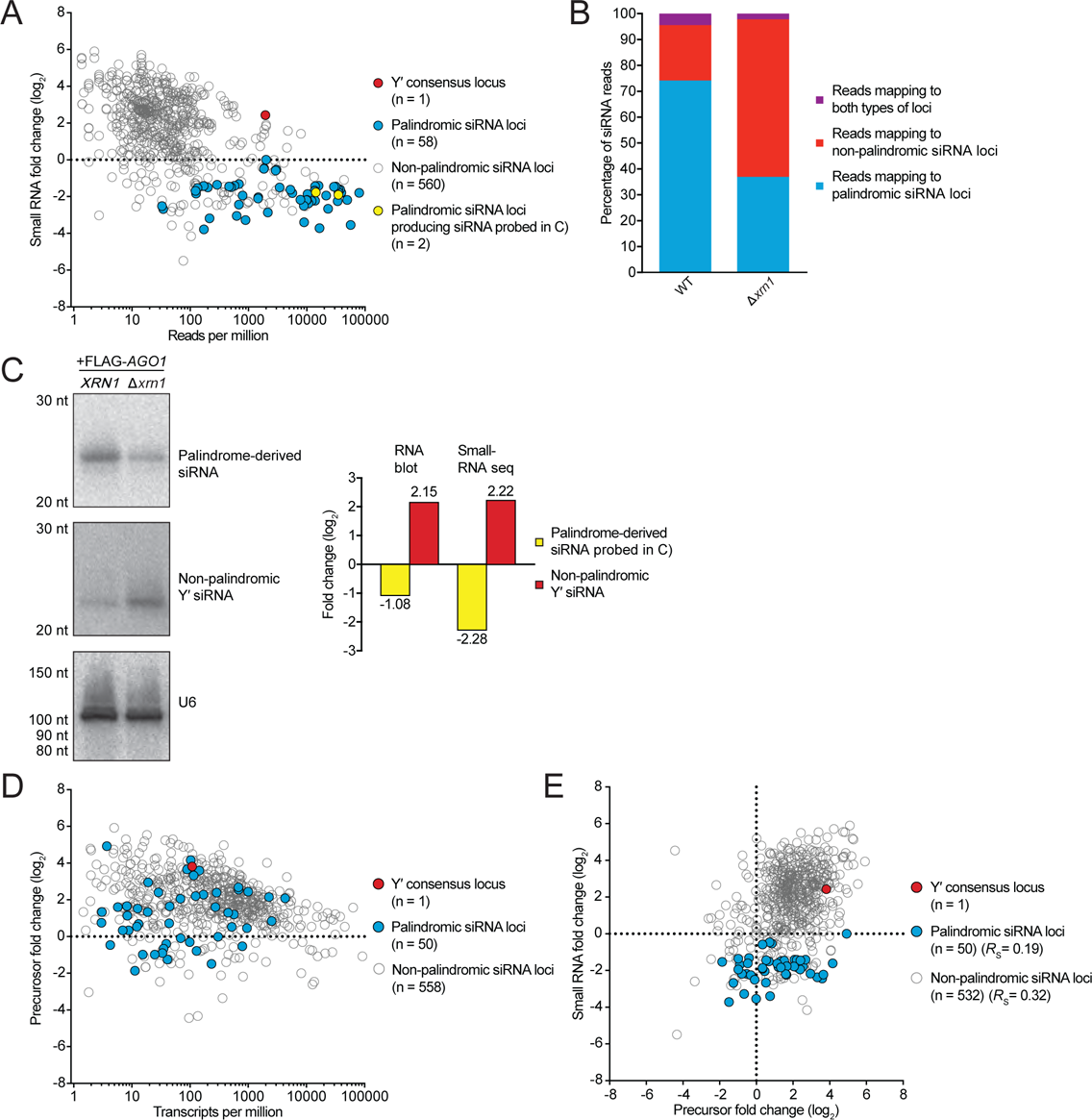
The effect of Xrn1p on the abundance of small RNAs and their precursors in *N. castellii*. **A)** The effect of Xrn1p on the abundance of small RNAs originating from siRNA-producing loci yielding at least 1 RPM (read per million) in WT cells. The scatter plot shows the fold change (log_2_) of 22–23-nt small RNAs deriving from each locus in the Δ*xrn1* versus WT strain (after normalizing to the recovery of internal standards) as a function of the abundance of 22–23-nt small RNAs deriving from the locus in the WT strain. The Y′ consensus locus is represented by the filled red circle, and other non-palindromic loci are represented by empty grey circles. The two palindromic loci probed in **C** are represented by the filled yellow circles, and the other palindromic loci are represented by the filled cyan circles. For each category, the number of loci is indicated in parentheses. Sequencing was from strains in which the endogenous Ago1p had been FLAG tagged. **B)** Overall effect of Xrn1p on the fraction of palindrome-derived and non-palindrome-derived small-RNA populations in *N. castellii*. The 22– 23-nt reads that mapped to siRNA-producing loci were designated as deriving from either palindromic loci (cyan), non-palindromic loci (red), or both types of loci (purple) in the same strains as **A**. **C)** Effect of Xrn1p on the expression levels of both a palindrome-derived siRNA and a non-palindrome-derived siRNA in *N. castellii*. At the left are phosphorimages of a small-RNA blot successively probed for siRNAs originating from either palindromic or non-palindromic precursor transcripts in the same WT and Δ*xrn1* strains as **A**. The migration of markers is indicated on the left. For a loading control, the blots were re-probed for the U6 snRNA. To the right of the blots is a bar graph showing the effect of deleting *XRN1* on these siRNAs, as measured both by RNA blots and by the sequencing in **A**. **D)** Effect of Xrn1p on the abundance of small-RNA precursor transcripts in *N. castellii*, as measured using RNA-seq. Shown for each precursor transcript expressed at ≥ 1 transcript per million (TPM) in WT cells is its fold change (log_2_) in the Δ*xrn1* versus WT strains (after normalizing to the recovery of internal standards) as a function of its abundance in WT cells. Strains were as in **A**, and precursor transcripts are classified as in **A**. **E)** Relationship between changes in small RNAs and changes in siRNA precursors observed after deleting *XRN1*. Shown for each precursor transcript expressed at ≥ 1 TPM in WT cells and that produced small RNAs at ≥ 1 RPM is the fold change (log_2_) in 22–23-nt small RNA reads observed in **A** as a function of the fold change (log_2_) in precursor transcripts observed in **D**. Representation of each type of locus as in **A**. The number of loci and Spearman’s rank correlation coefficient (*R*_s_) for each category is indicated in parentheses.

To ascertain if changes in siRNAs observed in the Δ*xrn1* strain could be attributed to changed accumulation of their siRNA-precursor transcripts, we performed RNA sequencing (RNA-seq) of longer transcripts, again mapping sequencing reads to annotated siRNA-producing loci. Many siRNA-precursor transcripts increased in expression in the Δ*xrn1* strain (Figure 3D, median change, 2.2- and 3.8-fold for palindromic and non-palindromic loci, respectively). Examination of the relationship between changes in siRNAs and changes in precursor transcripts uncovered a modest correlation for products of non-palindromic loci (*R*_S_ = 0.32; *P* < 0.001), with increased levels observed for both siRNAs and precursors from 430 of 532 loci (Figure 3E). In contrast, little correlation was observed for products of palindromic loci (*R*_S_ = 0.19; *P =* 0.2).

Similar expression changes of small RNAs and their precursor transcripts were observed when comparing the *N. castellii* selection strain and a Δ*xrn1* strain derived from it (Supplemental Figure 1B). In these strains, the most abundant siRNAs mapped to the loci expressing hairpins corresponding to the exogenous *HIS3*, *URA3,* and *GFP* genes. The siRNAs from these three loci decreased by 2.7-, 1.8- and 1.1-fold, respectively, in the Δ*xrn1* strain (Supplemental Figure 1B). The precursor transcripts for both the endogenous palindromic and non-palindromic siRNAs also increased in the Δ*xrn1* selection strain (Supplemental Figure 1C, median fold change of 3.5 and 4.6, respectively). As with the non-selection strains, when examining the relationship between the changes in siRNAs and the changes in precursor transcripts, a modest correlation was observed for products of non-palindromic loci (Supplemental Figure 1D, *R*_S_ = 0.33; *P* < 0.001), and little correlation was observed for products of palindromic loci (Supplemental Figure 1D, *R*_S_ = 0.04; *P =* 0.38).

These results suggest that the less efficient silencing of *GFP* observed in the Δ*xrn1* strain was likely at least partially attributable to reduced siRNAs deriving from the palindromic transcripts designed to silence *GFP*. One possibility is that the loss of *XRN1* causes an increase of non-palindromic siRNA precursors, and the increased siRNAs deriving from these precursors compete with the palindrome-derived siRNAs for loading into Ago1p, thereby causing more of the *GFP* siRNAs to be degraded before loading into Ago1p. In this scenario, Xrn1p could influence the levels of non-palindromic siRNA precursors either directly, by degrading these precursors, or indirectly, by increasing the efficiency of RNAi-mediated degradation of these transcripts, which are targets of as well as precursors for siRNAs of the RNAi pathway.

### Xrn1p enhances RNAi in vitro

To study the ability of Xrn1p to enhance RNAi, we purified wild-type *N. castellii* Ago1p and wild-type and catalytically impaired versions of *N. castellii* Xrn1p (each from *S. cerevisiae* overexpression strains that lacked *XRN1*) and examined the influence of Xrn1p activity on passenger-strand cleavage and target slicing in vitro. The in vitro assay used an siRNA duplex with a 5′-labeled passenger strand and a cap-labeled target RNA to follow both passenger-strand cleavage and target slicing in multiple-turnover conditions (Figure 4A). In the presence of wild-type (WT) but not catalytically impaired (mut.) Xrn1p, the amount of passenger-strand cleavage fragment substantially increased at early time points and then dropped at later time points (Figure 4B–C). The reduced passenger-strand fragment observed at later time points indicated that Xrn1p degraded this fragment, either as it exited RISC or after it dissociated from RISC—in either case, potentially enhancing RNAi efficiency by facilitating RISC maturation or preventing cleaved passenger strand from inhibiting RISC activity, respectively. An siRNA duplex with a 3′-labeled passenger strand showed a similar trend as the siRNA duplex with a 5′-labeled passenger strand; in the reaction with wild-type Xrn1p, the 3′-cleavage fragment of the passenger strand accumulated to a greater extent at early time points and then was degraded at later time points (Supplemental Figure 2A–C). These results indicated that Xrn1p degrades both cleavage fragments of the passenger strand in vitro—an observation consistent with the known preference of Xrn1p for substrates with a 5′ monophosphate.

**Figure 4.**
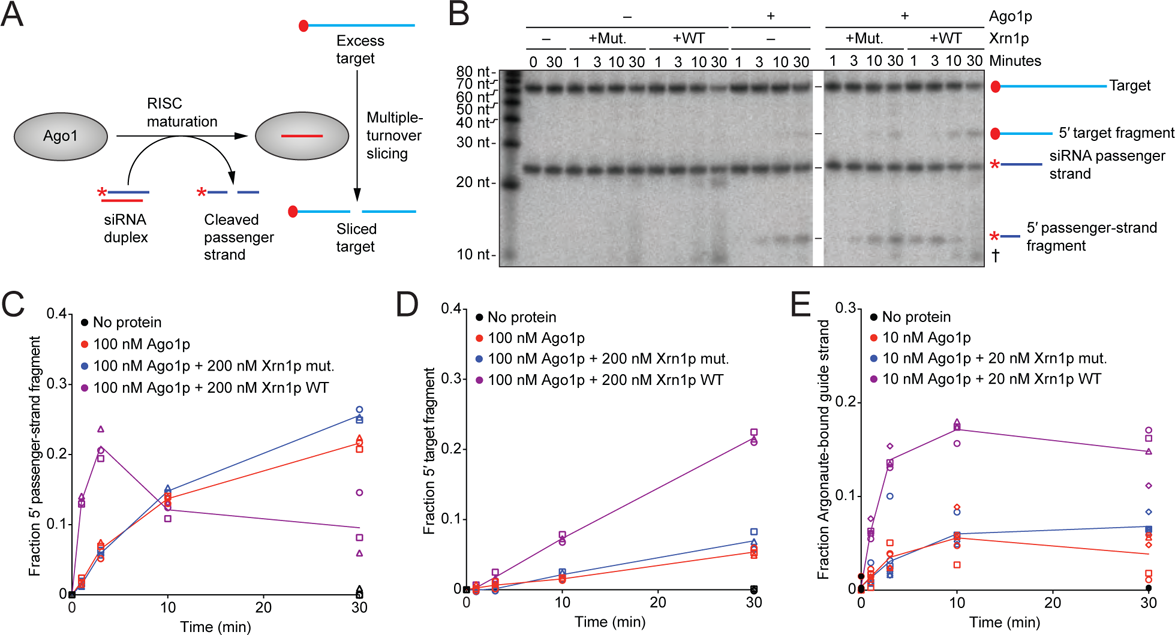
The impact of Xrn1p on siRNA loading, passenger-strand degradation, and target slicing. **A)** Experimental scheme of a combined assay monitoring both passenger-strand cleavage and target slicing. An siRNA duplex is loaded into Ago1p, which cleaves and discards the passenger strand before beginning to catalyze multiple-turnover slicing of the target RNA. Initial concentrations of siRNA duplex and target RNA were 10 and 100 nM, respectively. In the duplex, the red star indicates radiolabeled monophosphate on the 5′-end of the passenger strand. On the target, the red-filled circle indicates a 5′-radiolabeled cap. **B)** The effect of Xrn1p on Ago1p-catalyzed passenger-strand cleavage and target slicing, in the assay schematized in **A**. Shown is a representative denaturing gel that resolved cap-labeled target, cap-labeled target fragment, 5′ end-labeled passenger strand and 5′ end-labeled passenger-strand fragment after incubation for the indicated time, with or without purified Ago1p (100 nM) and with or without purified Xrn1p (WT or mutant [mut.]) (200 nM). The cross symbol (^†^) indicates degradation of the target after adding Xrn1p, which was observed at late time points and presumed to be due to a contaminant in the Xrn1p preparation. **C)** Quantification of the passenger-strand cleavage. Results are shown for three replicates (circles, squares, triangles) of reactions with either no protein (black), Ago1p only (red), Ago1p with Xrn1p mut. (blue), or Ago1p with Xrn1p WT (purple). The fraction of passenger-strand fragment was calculated by dividing the signal of product / (product + substrate). The lines, added for clarity, connect mean values and are not fit to an equation. **D)** Quantification of the 5′ target fragment. Otherwise, as in **C**. **E)** The effect of Xrn1p on siRNA association with Ago1p. Shown are results of a filter-binding assay that measured association of 5′ radiolabeled guide RNA with Ago1p after incubating components of the combined assay of passenger-strand cleavage and target slicing. Each assay included siRNA duplex, unlabeled target RNA, and the other components indicated, each at the same concentrations used in **B**. Results are shown for five independent experiments (circles, squares, triangles, diamonds, hexagons), which together contributed at least three data points for each time point for reactions with either no protein (black filled circles), Ago1p only (red), Ago1p and Xrn1p mut. (blue), or Ago1p and Xrn1p WT (purple). The fraction of Ago1p-bound guide strand was calculated by dividing the signal for the RNA that bound to the nitrocellulose membrane by that of the RNA that bound to the nylon and nitrocellulose membranes combined.

Wild-type Xrn1p also increased the rate of target slicing (Figure 4D). The increased rates of both passenger-strand cleavage and target slicing suggested that Xrn1p increased the amount of Ago1p that could be loaded with an siRNA duplex, perhaps by degrading non-specifically bound cellular RNAs that might be occluding siRNA loading. To assess whether Xrn1p activity increased the amount of Ago1p that could be loaded with a siRNA duplex, we performed filter-binding assays to measure the amount of siRNA bound to protein after incubating guide-labeled siRNA duplex with the purified proteins (Figure 4E). Preincubation of wild-type Xrn1p with Ago1p enhanced the amount of siRNA bound to protein, whereas preincubation with mutant Xrn1p did not have this effect (Figure 4E). These results supported the idea that the exonuclease activity of Xrn1p might help eliminate non-siRNA species that spuriously bind Ago1p and competitively inhibit siRNA duplex binding.

### *XRN1* affects the stability of the passenger strand in vivo

Because *N. castellii* Xrn1p degraded the cleaved passenger strand in vitro, we asked if it had this effect in vivo. We identified perfectly paired guide–passenger duplexes for which 22–23-nt reads were observed among small-RNA reads from wild-type, Δ*xrn1*, and *AGO1* slicing-impaired (*ago1*-*D1247R*) *N. castellii* strains. For each of these duplexes, the strand with more reads in the wild-type strain was designated as the guide, the other strand was designated as the passenger, and duplexes for which the passenger-to-guide ratio increased in the *ago1* mutant strain were carried forward as experimentally supported siRNA duplexes. Twelve-nucleotide reads that perfectly matched the 3′ end of the passenger strand were designated as 3′-cleavage fragments of passenger strands, and depending on whether the duplex was comprised of 22- or 23-nt species, 10- or 11-nt reads that perfectly matched the 5′ end of the passenger strand were respectively designated as 5′-cleavage fragments of passenger strands. For most duplexes, disrupting *XRN1* increased the ratio of full-length passenger strand to guide strand, as would be expected if Xrn1p facilitates RISC loading (Figure 5A, median increase, 2.3-fold; *P* < 0.001, Wilcoxon signed-rank test). Disrupting *XRN1* also increased the ratio of cleaved passenger strand to guide strand modestly (Figure 5B, median increase, 1.3-fold; *P* < 0.001).

**Figure 5.**
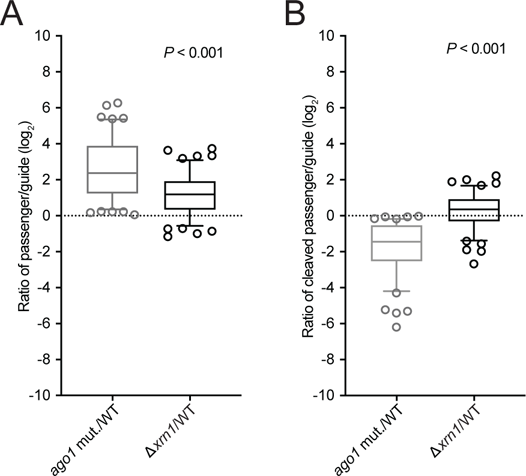
The influence of Xrn1p on the accumulation of the siRNA passenger strand and its cleavage fragments in vivo. **A)** Accumulation of full-length passenger strands. Box-and-whisker plots show the log_2_ ratios of the reads for the full-length passenger strand over the reads for the full-length guide strand in the slicing-impaired *ago1* mutant (mut.) versus WT strains and in Δ*xrn1* versus WT strains. Results from 114 guide–passenger duplexes are plotted, with the line indicating the median, the box indicating 25–75 percentiles, the whiskers indicating the 5–95 percentiles, and open circles indicating individual outliers. The two-tailed Wilcoxon signed-ranked test was used to evaluate the similarity of the two distributions (*P* value). **B)** Accumulation of cleaved passenger strands. Box-and-whisker plots show the log_2_ ratios of the reads for the cleaved passenger strand over the reads for the full-length guide strand in *ago1* mut. versus WT strains and in Δ*xrn1* versus WT strains. Otherwise, as in **A**.

We also examined the length distributions and the first nucleotide composition of the genomically mapping 9–26-nt reads in the libraries of each of these strains, excluding reads that mapped exclusively to rRNAs or tRNAs (Supplemental Figure 3). There were no substantial differences in the size distributions of the 9–26-nt reads in the Δ*xrn1* or *AGO1* slicing-impaired strains compared with the wild-type strain, although there were slightly more 15–20-nt reads in the Δ*xrn1* strain (Supplemental Figure 3). Similar to previous studies, most 23-nt RNAs began with U, which suggested that most siRNA guide strands begin with this nucleotide (8,54). However, in the Δ*xrn1* and *AGO1* slicing-impaired strains a higher fraction of 23-nt reads began with non-U nucleotides (48.6 and 51.3% respectively, compared to 47.6% for the wild-type strain), which suggested that full-length passenger strands were stabilized in these mutant strains (Supplemental Figure 3). Additionally, the Δ*xrn1* and *AGO1* slicing-impaired strains also had a higher fraction of 23-nt reads with A in the third to last position (24.8 and 36.0% respectively, compared to 19.3% for the wild-type strain), which would be expected of full-length passenger strands that could pair with guide strands that begin with U. These results indicated that Xrn1p affects the stability of full-length and cleaved passenger strands in vivo, as it does in vitro.

### Xrn1p enhances target slicing and eliminates the 3′ slicing fragment

To learn whether Xrn1p enhances the efficiency of any steps following duplex loading, RISC maturation, and passenger-strand degradation, we purified mature *N. castellii* RISC programmed with a specific guide RNA (55) and examined the ability of *N. castellii* Xrn1p to enhance slicing of cap-labeled target in vitro. Under single-turnover conditions, in which the RISC enzyme was in excess over target, Xrn1p activity had no detectable influence on slicing rates (Supplemental Figure 4A–B). Likewise, Xrn1p activity had no detectable influence on the initial burst phase of a multiple-turnover slicing reaction (Figure 6A), in which target was in 10-fold excess over the RISC enzyme (Figure 6B–C). However, after the slicing product reached the concentration of RISC (10 nM) and the reaction entered a slower phase characterized by rate-limiting product release, wild-type Xrn1p but not the catalytically impaired Xrn1p increased the rate of target slicing (Figure 6B–C). Enhancement of this second phase of target slicing increased after boosting the Xrn1p concentration from 10 to 30 nM but did not increase with even more Xrn1p (Figure 6C).

**Figure 6.**
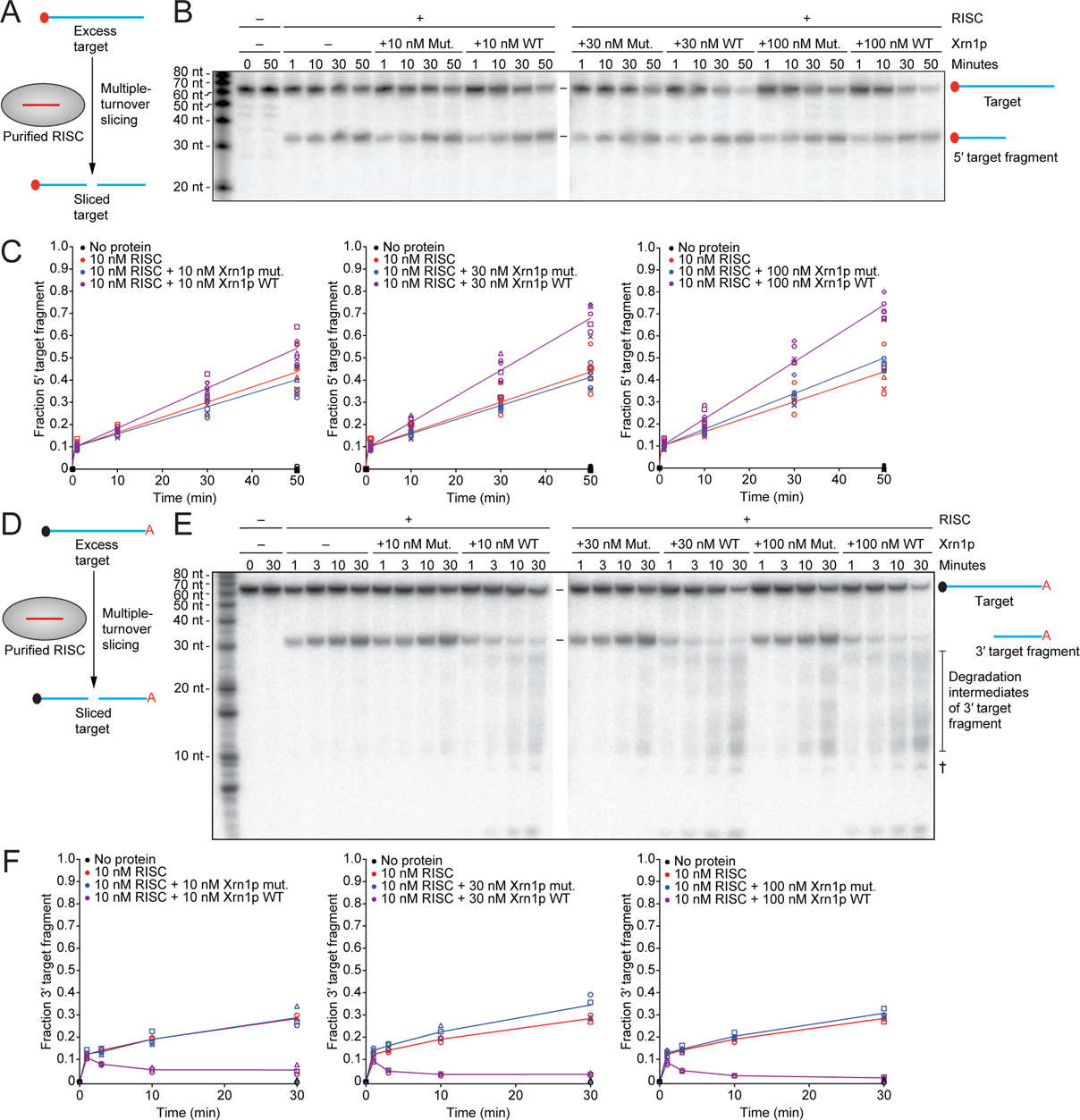
The impact of Xrn1p on multiple-turnover slicing and on stability of the 3′ target fragment. **A)** Experimental scheme of the assay for multiple-turnover slicing using a cap-labeled target. The red-filled circle on the target RNA indicates a radiolabeled 5′ cap. The initial concentration of target was 100 nM. **B)** The effect of Xrn1p on multiple-turnover slicing in the assay schematized in **A**. Shown is a representative denaturing gel resolving target from its sliced product after incubation with or without purified RISC–miR-20a (10 nM), with the indicated concentration of Xrn1p (WT or mut.; 0, 10, 30, or 100 nM) for the indicated amount of time. **C)** Quantification of the 5′ product of target slicing. Results are shown for six independent experiments (circles, squares, triangles, diamonds, hexagons, and exes) of reactions with either no protein (black), RISC–miR-20a only (red), RISC–miR-20a with Xrn1p mut. (blue), or RISC– miR-20a with Xrn1p WT (purple). The fraction of 5′ target fragment was calculated by dividing the signal of product / (product + substrate). The lines indicate the best fit to the burst equation. **D)** Experimental scheme of the assay for multiple-turnover slicing with a 5′-capped, 3′-radiolabeled target. The black-filled circle on the target indicates a 5′-cap, and the red A indicates a radiolabeled cordycepin at the 3′-end of the target. The initial concentration of target was 100 nM. **E)** The effect of Xrn1p on multiple-turnover slicing in the assay schematized in **D**. The cross symbol (^†^) indicates degradation of the target after adding Xrn1p, which was observed at late time points and presumed to be due to a contaminant in the Xrn1p preparation. Otherwise, as in **B**. **F)** Quantification of the 3′ product of target slicing. Results are shown for three independent experiments (circles, squares, triangles). The lines, shown for clarity, connect mean values and are not fit to an equation. Otherwise, as in **C**.

These results suggested that Xrn1p enhances release of sliced product, presumably by degrading the 3′ product of slicing, which contains the 5′-monophosphate characteristic of Xrn1p substrates. To examine whether Xrn1p has this function, we performed a slicing reaction with a 5′-capped, 3′-labeled target RNA in multiple turnover conditions (Figure 6D). In reactions lacking Xrn1p or containing catalytically impaired Xrn1p, the 3′ slicing product accumulated rapidly during the burst phase and then more slowly during the part of the reaction characterized by rate-limiting product release, as observed for the 5′ product in reactions with cap-labeled target (Figure 6E–F). In reactions containing wild-type Xrn1p, the 3′ slicing product similarly accumulated at an early time point but then began to decrease as it was degraded by Xrn1p (Figure 6E–F). This rate of degradation increased as the concentration of Xrn1p increased from 10 to 30 nM but did not significantly increase with additional Xrn1p (Figure 6F). The elimination of this 3′ product of slicing, which pairs to the guide-RNA seed region, which in turn is critical for target recognition and binding (56-58), would prevent this 3′ slicing product from rebinding to the guide and would thereby allow RISC to bind and slice another target RNA.

In support of the proposal that Xrn1p degrades 3′ products of slicing, comparison of RNA-seq data from the selection strain with and without *XRN1* showed that, in the absence of Xrn1p, RNA preferentially accumulated in the mRNA regions downstream of *HIS3*, *URA3,* and *GFP* slicing (Supplemental Figure 4C). In contrast, endogenous genes that were not targets of RNAi in the selection strain showed a uniform increase in RNA sequencing reads mapping across the gene in the *XRN1* knockout strain, and showed an increased proportion of reads mapping to their 3′ ends in both WT and *XRN1*-deficient backgrounds (Supplemental Figure 4D). Thus, as observed in other species (29-31), *XRN1* is responsible for eliminating the 3′ products of slicing in budding yeast.

## Discussion

Our results show that Xrn1p influences most steps of the RNAi pathway in budding yeast (Figure 7). Xrn1p increases the levels of siRNAs that accumulate from palindromic loci, presumably by reducing the amount of non-palindromic siRNA-precursor transcripts, thereby reducing competition for entry into the pathway from siRNAs that would otherwise be produced from these non-palindromic precursors. Once the siRNA duplexes are produced, Xrn1p facilitates their loading into Argonaute, perhaps by eliminating other RNA species that spuriously bind to Argonaute and thereby inhibit duplex loading. After duplexes bind to Argonaute and passenger strands are cleaved, Xrn1p helps degrade the cleavage fragments, which prevents inhibition of RISC and might facilitate its maturation. Finally, Xrn1p facilitates multiple turnover of RISC by eliminating the 3′ products of slicing, thereby preventing product inhibition and perhaps helping to regenerate unbound RISC. We also detected a physical interaction between Ago1p and Xrn1p, which might play a role in one or more of these functions.

**Figure 7.**
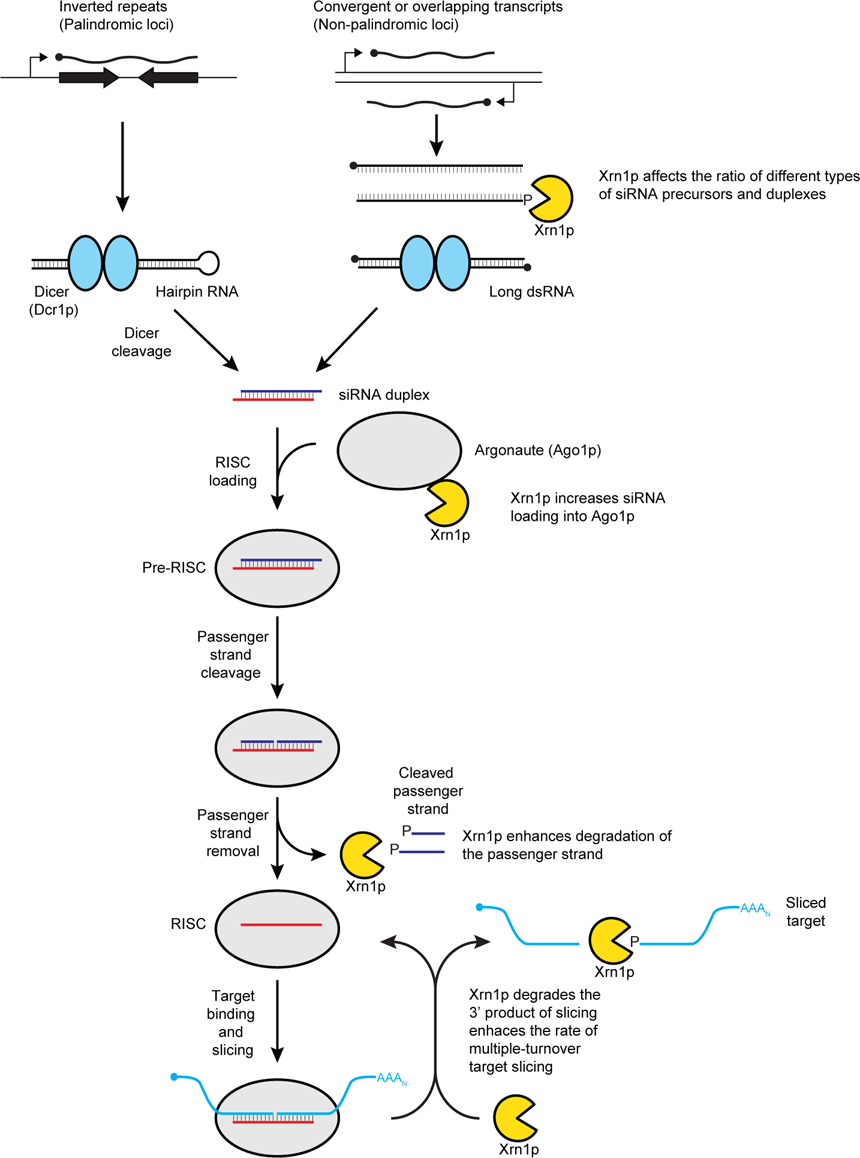
Schematic depicting the roles of Xrn1p in budding yeast RNAi. See main text for description.

Even with these new-found roles of Xrn1p, the RNAi pathway of budding yeast appears to be the most streamlined of any characterized to date, with only three known components: Dcr1p, Ago1p, and Xrn1p. Although we cannot rule out participation of other factors that might have been missed by our genetic and pulldown approaches, the notion that RNAi in budding yeast relies on only three proteins seems plausible. For example, the most notable difference between budding-yeast Ago1p and Argonaute proteins characterized from other lineages is that purified Ago1p can autonomously load an siRNA duplex and then cleave and remove the passenger strand of this duplex to form active RISC capable of slicing target RNA (59), whereas other Argonaute proteins require the Hsc70/Hsp90 chaperone and ATP to assume a conformation capable of duplex loading and RISC maturation (23,18,20-22). That Ago1p can autonomously perform these functions in vitro, with some enhancement by Xrn1p, suggests that other factors might not be required for duplex loading and RISC maturation in budding yeast.

The unusual domain structure and activity of budding-yeast Dcr1p might also obviate the need for other accessory factors needed for siRNA loading into Argonaute in other systems. Budding-yeast Dcr1p possesses only one RNase III domain, whereas canonical Dicers possess two RNase III domains (60). Consistent with RNase III domains always functioning in pairs (61), budding-yeast Dicer forms a homodimer to cleave both strands of the dsRNA (60). Whereas canonical Dicers measure from the end of the dsRNA to determine the site of cleavage (62,63), multiple Dcr1p homodimers bind cooperatively within the dsRNA, with the distance between active sites of adjacent homodimers determining the length of the siRNA duplexes (60). Each unit of the budding-yeast homodimer possesses two dsRNA-binding domains (dsRBDs), which are both required for accumulation of siRNAs to normal levels in vivo, although the second dsRBD is dispensable for producing siRNAs in vitro (60). One proposal is that in cells this second dsRBD acts after siRNA production, in tandem with the second dsRBD contributed by the other Dcr1p of the homodimer, to perform functions normally provided by dsRBD-containing Dicer accessory factors, such as helping to transfer the siRNA duplex from Dicer to Argonaute, thereby obviating the need for these cofactors (60).

Further supporting the existence of a streamlined pathway in budding yeast, *N. castellii* lacks orthologs of other factors that participate in RNAi and related pathways in other species. For example, *N. castellii* and all other RNAi-possessing budding-yeast species lack recognizable homologs of RdRPs (8). Budding yeast also lack a discernible ortholog of *HEN1*, which methylates the 2′ oxygen of the 3′-terminal nucleotide of plant and metazoan siRNAs, plant miRNAs, as well as metazoan Piwi-interacting RNAs (piRNAs), thereby protecting these small RNAs from 3′-end uridylation and degradation (64-74). Indeed, we found that *N. castellii* siRNAs are susceptible to periodate oxidation and beta-elimination, thereby confirming that they are not methylated (Supplemental Figure 5A).

*N. castellii* does have orthologues of several other factors thought to participate in RNAi in other species, including the QIP exonuclease, which removes the cleaved passenger strand in *Neurospora crassa* (26), and the autoantigen La RNA-binding protein, which is reported to enhance the multiple-turnover of RISC in Drosophila RNAi (75). However, disruption of the orthologs of the genes encoding QIP (*GFD2* and its paralog, *NCAS0A00350*), or the ortholog of La (*LHP1*) in an *N. castellii* RNAi-reporter strain had little effect on *GFP* silencing (Supplemental Figure 5B). Taken together our results suggest that Xrn1p helps degrade the cleaved passenger strand and enhances the multiple turnover of RISC in budding yeast, thereby obviating the need for other proteins to perform these functions.

The general eukaryotic mRNA decay factors *DCP2* and *SKI3* have also been implicated in small-RNA–mediated gene-silencing pathways. *DCP2*, which encodes the mRNA decapping enzyme, is involved in miRNA-mediated silencing but not RNAi in Drosophila (76). *SKI3*, which encodes a component of the Ski complex, enables the cytoplasmic exosome to degrade 5′-target-slicing fragments in Drosophila (30). Although disrupting *DCP2* and *SKI3* affected the abundances of siRNAs in the cell (Supplemental Figure 1A), we found that the deletions of these genes in an *N. castellii* RNAi-reporter strain had little-to-no effect on *GFP* silencing (Figure 1E). These results suggest that each protein is either not involved in budding-yeast RNAi or has effects on the pathway that are undetectable in the *GFP*-silencing assay.

Xrn1p satisfies the expected criteria of a cofactor participating in the budding-yeast RNAi pathway. It is present in all known eukaryotes, playing important roles in RNA metabolism critical for cellular fitness, which explains why it was not lost in *S. cerevisiae* and other species that have lost RNAi. Although not thought to be an essential component of RNAi in any species, Xrn1 has been shown to function in RNAi and related pathways in diverse species, and our observations in budding yeast expand this repertoire, both with respect to its evolutionary reach and with respect to its functions within a single species. Indeed, studies in other species have focused on the role of Xrn1 on a particular step of the pathway, and thus it will be interesting to learn the degree to which Xrn1 has multiple functions in RNA-silencing pathways of other eukaryotes or whether these other lineages have evolved cofactors that assume these more specialized roles in more elaborated pathways.

## Acknowledgements

We thank Kathy Lin, Kathy Xie, George Bell, Prat Thiru, and Brian Chin for helpful discussions, and Katrin Heindl and Sean McGeary for assistance with the protein purifications.

## Contributions

M.A.G. and I.A.D. made the strains for the genetic selection. I.A.D. carried out the pilot genetic selection with help from M.A.G., and M.A.G. performed the genetic selection with modified strains and analyzed the genomic sequencing data. D.E.W. made the tagged Ago1p constructs, and carried out the immunoprecipitations and protein blots, the microscopy, and RNA blots. M.A.G. performed all small-RNA and RNA sequencing and analysis. M.A.G. purified *N. castellii* Ago1p and Xrn1p and did all of the biochemistry experiments and analysis. All authors contributed to experimental design. M.A.G. and D.P.B. wrote the manuscript, and all authors edited the manuscript.

## Methods

### Growth conditions and genetic manipulations

*N. castellii* was grown at 30°C on standard *S. cerevisiae* liquid and solid media (YPD and SC), unless otherwise indicated. Transformations were performed as described (1), with the following modifications: The transformation mix (cells + DNA + PEG + lithium acetate) was incubated at 23°C for 30 min with rotation and without DMSO. The mixture was then incubated at 37°C for 20 min, resuspended, and plated on selective media.

### Plasmid construction

A list of plasmids generated in this study is provided (Table S6).

#### Integrating plasmids for N. castellii RNAi selection strains

pIp-strongSC_GFP, the integrating plasmid containing the *GFP*-silencing construct, was constructed previously (1). To construct pRS403-ScerURA3hp-HIS3-PEST, the integrating plasmid containing the *URA3*-silencing construct, the N. castellii genomic sequence downstream of the ORF NCAS0C02950 was amplified (primers: Ncas_Int679_For 5′ GGCAAATTTGTATGAGGGATAAA and Ncas_Int679_Rev 5′ TAATTCGATTACGTTAGCTGTT) and cloned into pRS403-pGAL1-hpSC_URA3 (1) using PsiI and NaeI restriction enzymes (New England Biolabs, NEB). In addition, the *PEST* sequence from S. cerevisiae CLN2 was appended to the C-terminal codon of the HIS3 gene using the In-Fusion Cloning Kit (Clontech) as previously described (2), which increased the stringency of the selection. To construct pRS405-ScerHIS3hp, the integrating plasmid containing the HIS3-silencing construct, a 286 bp of HIS3 sequence from pRS403-pGAL1-hpSC_URA3 was amplified (primers: HIS3_hp_For 5′ AAAAGCTTGACCGAGAGCAA and HIS3_hp_Rev 5′ GCGTATTACAAATGAAACCAAGATTCA) and initially cloned into pRS405 (3) between HindIII and XhoI (NEB). A DNA segment containing the *GAL1* promoter, the *HIS3* sequence in the antisense direction, and the Schizosaccharomyces pombe rad9 intron was synthesized via fusion PCR (4) and inserted between BamHI and PstI (NEB). The N. castellii genomic sequence in between the ORFs NCAS0D00680 and NCAS0D00690 was amplified (primers: Ncas_Int618_For 5′-GTTCGCCGGCCTTCCCGCGCTATGAAATTA and Ncas_Int618_Rev 5′-ATCAGGCGCCGAGCATAACCGCTCAAATGC) and inserted between the NaeI and KasI (NEB) restriction sites. To construct pRS404-NcasAGO1, the integrating plasmid containing N. castellii AGO1 (NCAS0J02110), N. castellii genomic sequence downstream of the ORF NCAS0C00690 was amplified (primers: Ncas_Int696_For: 5′-GGCCGGTACCAATTCATCTAGCAGGATGTAAAATG; Ncas_Int696_Rev: 5′-GAAAGCCGGCGTAGAGCATGCGAGGTTTGG) and inserted between the KpnI and NaeI (NEB) restriction sites in pRS404 (3). The *AGO1* ORF along with its upstream and downstream sequence was amplified (primers: AGO1-Intergenic-For 5′-GCTGGAGCTCTGAACGTGTGGAAGACCAAA; AGO1-Intergenic-Rev 5′-ATGACTCGAGAGTGGCTAACGGCAACATATC) and inserted between the SacI and XhoI (NEB) restriction sites of pRS404. To construct pRS402-NcasDCR1, the integrating plasmid containing N. castelii DCR1 (NCAS0C00230), N. castellii genomic sequence upstream of the ORF NCAS0E03540 was amplified (primers: Ncas_Int701_For 5′-ATTCGGATCCTGCAGGCTGTTTGCTGTACT; Ncas_Int701_Rev 5′-GGTGGCGGCCGCGGGGTAACTATCCGCGTCTAA) and inserted between the BamHI and NotI (NEB) restriction sites in pRS402 (5). The *DCR1* ORF along with its upstream and downstream sequence was amplified (primers: DCR1-Intergenic-For 5′-CCCCCTCGAGTTTGTAAAGAAATTGATGCTTCG; DCR1-Intergenic-Rev 5′-TGCAGGATCCGAATCTGGTATGGGATCATATTGG) and inserted between the XhoI and BamHI (NEB) restriction sites of pRS402.

#### Plasmids for protein expression strains

pYES2.1-FLAG3-SUMO-TEV-NcasAGO1, which expresses N-terminally FLAG- and SUMO-tagged *AGO1*, was constructed by creating a gBlock gene fragment (IDT) of 3x-FLAG, SUMO, and TEV sequences and cloning it immediately downstream of the *GAL1* promoter in pYES2.1 by Gibson Assembly (NEB). The *AGO1* CDS lacking the start codon was then inserted immediately downstream of the 3x-FLAG-SUMO-TEV construct by Gibson Assembly. pYES2.1-NcasXRN1-TEV-SUMO-FLAG3, which expresses C-terminally FLAG- and SUMO-tagged *XRN1* was constructed in the same manner, except the full *XRN1* CDS was inserted in between the *GAL1* promoter and the TEV-SUMO-3x-FLAG sequence. QuikChange Site-Directed Mutagenesis (Agilent) was used to generate pYES2.1-NcasXRN1(D206N-D208N)-TEV-SUMO-FLAG3, for which active-site aspartates at amino acid positions 206 and 208 of *XRN1* were substituted with asparagine (QuikChange primers, with changed nucleotides bolded: sense 5′-GAAGAATAACAGCAGAGAAGGATTTTGATGAAAATACTAGACACTGTATTTATGGA TTA**A**ATGCA**A**ATCTAATAATATTGGGGCTTTCCACTCATGCTCCACATTTCGCACTGTTAAGAG; antisense, reverse complement of this sequence).

### Strain construction

A list of strains generated and used in this study is provided (Table S5).

#### N. castellii RNAi selection strains, RNAi mutant strains and derivatives

The parental *N. castellii* genetic selection strains DPB537 (*MAT* **a**) and DPB507 (*MAT* ⍺) were derived from the original progenitor strains DPB004 and DPB005 (1), respectively. *ADE2* (NCAS0H02150), *HIS3* (NCAS0B01740), *LEU2* (NCAS0C04870) in both strains, as well as *LYS2* (NCAS0I01040) in DPB537 and *MET17* (NCAS0A05810) in DPB507, were disrupted with loxP-kanMX6-loxP modules of plasmid pUG6 (6) that had been fused via PCR (7) with ∼400–500-bp targeting arms on both sides of the cassette followed by expression of Cre recombinase, as described (1). *TRP1* (NCAS0E01400) was disrupted using the hygromycin cassette of pAG32 (8), as described (1). The GFP(S65T)-kanMX6 module was inserted into the genome as described (1). The *GFP* silencing construct (pIp-strongSC_GFP) was integrated upstream of the *ADH2* ORF (NCAS0I02350), as described (1) in these strains and all strains containing this *GFP* silencing construct. The integrating plasmids generated from the pRS402, pRS403, pRS404, and pRS405 backbones were each digested with restriction enzymes that made single cuts in their regions of homology with the *N. castellii* genome and then transformed into *N. castellii* independently. Transformants were selected on appropriate synthetic media based on the selectable markers they possessed. Strain DPB325 was derived from strain DPB313, which was previously constructed (1), by disrupting *ADE1* (NCAS0I01850) with a cassette consisting of ∼400–500-bp targeting arms upstream and downstream of the *ADE1* ORF amplified from genomic DNA attached to the kanMX cassette of pFA6a via fusion PCR. Strain DPB079 was derived from strain DPB005 using the loxP-kanMX6-loxP modules discussed above to disrupt *HIS3* and *MET17*. DPB534 was created by amplifying regions upstream and downstream of *DCR1* (primers: Upstream arm: 5′-TAATTAGCGGGCCATTATACCG; 5′-TTGTATAATTGCGTGTAGGTCACTT; Downstream arm: 5′-AAATGATATTTATGCACCTTTT; 5′-TTGAAGTGGAAGCCAGAATG) and appending them to a *NAT* cassette from pAG25 (8) via fusion PCR. The *XRN1* (NCAS0C04170) disruption cassette was created by amplifying genomic regions flanking the *XRN1* ORF (primers: Upstream arm: 5′-TTGCTCTCGAGCTTAGCAAC; 5′-TGGAATACCCATGATTAAAATGAA; Downstream arm: 5′-ATGCCACCACCACATCCATA; 5′-AGATGGGATGGCGTTGATAG) and adding them to a *NAT* cassette via fusion PCR as described above. Strains DPB510, DPB535, and DPB541 were created by transforming the *XRN1-NAT* disruption cassette into strains DPB507, DPB079, and DPB537, respectively, and transformed and selected on YPD + NAT solid media. Similarly, the *SRB8* (NCAS0D04810) disruption cassettes were created by amplifying flanking genomic regions (Upstream arm: 5′-AGGGGAAGAATGTTTTATTTGTTT; 5′-AAAAAGTTGGGGGTTCTCTAGT; Downstream arm: 5′-GCACGACGATTTTGAGACAA; 5′-CCATCTTCAAAGAGGCCCCATA) and adding them to a *NAT* cassette or a *N. castellii LYS2* cassette consisting of the ORF with upstream and downstream regions (5′-ACCAATCGACCTTTGGATTA; 5′-TGATTCGGTTTTACAAATTGTTC) by fusion PCR. Strains DPB578 and DPB582 were created by transforming the *SRB8-NAT* or *SRB8-LYS2* followed by *XRN1-NAT* disruption cassettes, respectively, into strain DPB537 and then selected as discussed above.

#### RNA decay mutants

The RNA decay mutant strains DPB334, DPB336, DPB337 and DPB339 were generated from the parental strains DPB331 and DPB333 (1). AGO1 was disrupted in strain DPB442 with a hygromycin cassette as previously described (1). Flanking regions of DCR1 (primers: Upstream arm: 5′-ACTGATCCGGAAAAGAGAGT; 5′-TCACTTGATCTGTTGCTGGAGG; Downstream arm: 5′-AGGCATTGCAACAATCTGTGA; 5′-GAGTTTATCGATGATACCATTGAAGG), XRN1 (primers: Upstream arm: 5′-TTGCTCTCGAGCTTAGCAAC; 5′-TGGAATACCCATGATTAAAATGAA; Downstream arm: 5′-ATGCCAATGCCACCACCACA; 5′-AGATGGGATGGCGTTGATAG), SKI3 (NCAS0A15300 – primers: Upstream arm: 5′-TTGCACTGGTTGACTCCTCT; 5′-CAATTGCTTCACTTCCGACA; Downstream arm: 5′-GGCTGTTATGGCTTTGAAGG; 5′-TCAAAATAGGCTTATCCGACGA) and DCP2 (NCAS0G02500 – primers: Upstream arm: 5′-ATATGGCAGGCGTGTTTGGT; 5′-AGGGTGTGACAGAGGAAAAGT; Downstream arm: 5′-GGTTACGATCTATTGCTGTCATTC; 5′-CCATCAAACTCACACATTCAAAA) were amplified from genomic DNA. These flanking arms were appended to the hygromycin cassette of plasmid pAG32 via fusion PCR. These disruption cassettes were transformed into strains DPB331 and DPB333 and transformants were selected for on YPD + hygromycin solid media.

#### Tagged Ago1p, Dcp2p, and Xrn1p strains

Diploid strain DPB202 was made by transforming the previously synthesized AGO1 hygromycin disruption cassette into the diploid strain Y235 (1) and selecting for transformants on YPD + hygromycin solid media. An N-terminal-kanMX-pGAL1-3x-HA-AGO1 tagging construct was generated by amplifying a region of the pFA6a-kanMX6-pGAL1-3HA plasmid (9) and joining it via fusion PCR to ∼400–500-bp targeting arms corresponding to the regions immediately upstream of the AGO1 CDS and immediately downstream of the AGO1 start codon. To create diploid strain DPB201, this AGO1 tagging construct was transformed into diploid strain DPB202 and transformants were selected for on YPD + G418 solid media. In strains DPB215 and DPB221, XRN1 was C-terminally tagged with 3x-HA and kanMX was created via fusion PCR (7). Genomically encoded, N-terminally tagged eGFP(S65T)-AGO1 was created by replacing the start codon of AGO1 with S. cerevisiae URA3 expression cassette amplified from pYES2.1 (Invitrogen) (1) and appended to the AGO1 arms of homology described above. These cassettes were transformed into strain DPB211 and transformants were selected on SD – Ura. The eGFP(S65T) construct was amplified from strain DPB314 (1) and transformed into the intermediate strain containing the URA3 cassette in place of the AGO1 start codon and transformants were selected on media containing 5-FOA. In strains DPB231 and DPB234, XRN1 and DCP2 were C-terminally tagged with cassettes containing mCherry and kanMX for selection from pBS34 (10) with fusion PCR.

#### Strains for small-RNA and RNA sequencing

DPB220 served as the parental strain for all strains used for small-RNA and RNA sequencing. To create DPB228, the AGO1 slicing-mutant strain, the sequence encoding the PIWI domain of AGO1 (NCAS0J02110 - Ncas_Chr10: 404375–405853) was replaced by transforming with S. cerevisiae URA3 expression cassette amplified from pYES2.1 flanked by ∼400–500-bp arms of homology, as described before. QuikChange Site-Directed Mutagenesis (Agilent) was used to change AGO1 coding sequence in pYES2.1-AGO1 (1) to encode arginine in place of aspartate at amino acid position 1247 (D1247R) (QuikChange primers, changed codon is in bold: 5′-GTCGCATCTCCTGTTTATTACGCTCGTTTATTGTGTGAACGTGGTGCTGCA; 5′-TGCAGCACCACGTTCACACAATAAACGAGCGTAATAAACAGGAGATGCGAC). The ago1 D1247R construct was amplified and transformed into the intermediate strain, and transformants were selected as described above. To create DPB622, the Δxrn1 strain, the flanking regions of *XRN1* previously used for the hygromycin disruption cassette were amplified from genomic DNA and appended to the kanMX cassette of pFA6a via fusion PCR. This disruption cassette was transformed into strain DPB220 and transformants were selected on YPD + G418.

#### Protein-expression strains

The *S. cerevisiae* protein-expression strains DPB597, DPB598, DPB1100, and DPB1101 were synthesized using the parental yRH101 strain (a gift from Stephen Bell, MIT), which was derived from ySC7 (11). To create strain DPB597, *XRN1* was deleted in yRH101 using a CRISPR–Cas9 system adapted for use in *S. cerevisiae* (12,13). The template for the sgRNA targeting *XRN1* (Genome matching sequence in bold. 5′-GATCG**AAAAGGTTTTAGGTTACGCGG**; 5′-**AAAACCGCGTAACCTAAAACCTTTT**C) was cloned into plasmid pV1382 and transformed into yRH101 along with a repair template made from hybridizing oligos with homology upstream and downstream of the *XRN1* ORF (5′-AAAAATCAACACTTGTAACAACAGCAGCAACAAATATATATCAGTACGGTAACATA CGAC; 5′-GATATACTATTAAAGTAACCTCGAATATACTTCGTTTTTAGTCGTATGTTACCGTACTGA). To create strains DPB598, DPB1100, and DPB1101, plasmids pYES2.1-FLAG3-SUMO-TEV-NcasAGO1, pYES2.1-NcasXRN1-TEV-SUMO-FLAG3, and pYES2.1-NcasXRN1(D206N-D208N)-TEV-SUMO-FLAG3, respectively, were transformed into strain DPB597, and transformants were selected on SD – Ura.

### EMS mutagenesis and selection

Mutagenesis of DPB537, the N. castellii genetic-selection strain, with ethyl methane sulfate (EMS) was carried out as described (14), except scaled-up tenfold. An overnight YPD culture of N. castellii was used to inoculate a large YPD (∼150 mL) culture that was grown until OD600 1.0, split equally into aliquots that contained ∼2.1 x 109 cells, one of which was incubated with 300μL EMS and the other which was incubated with 300 μL sodium phosphate buffer. The OD600 of each culture was measured and was used to calculate the dilutions needed to plate 100 mutagenized or control cells on separate YPD plates to determine the death rate of cells in the mutagenesis. Mutagenized cells were split into four equal aliquots and plated on SGal – His, – Leu, – Ura (drop-out media); SD – His, – Leu, – Ura (drop-out media); SM (synthetic minimal) + 2% galactose + Lys (SGal + Lys) (drop-in media); and SM (synthetic minimal) + 2% glucose + Lys (SD + Lys) (drop-in media) agar plates (245 mm x 245 mm). These plates were incubated at 25°C until colonies formed.

### Flow cytometry and mating of RNAi mutant strains

A culture of each mutant strain that had grown on selective media was inoculated in 200 μL inducing SC media (2% galactose) in a 96-well plate and grown to saturation. Fresh cultures were then inoculated at OD_600_ 0.2 with cells from the saturated cultures and were grown until all strains were in log phase (∼6 h). Cells were analyzed using FACSCalibur (BD Biosciences), and data were processed with CellQuest Pro (BD Biosciences) and FlowJo (Tree Star). For each experiment, a gate for eGFP fluorescence was defined using the WT genetic-selection strain (DPB537) as a negative control to define the boundaries of the gate. Strains for which > 0.5% of the population was in the eGFP-positive gate were carried forward for complementation analysis.

Matings of candidate *N. castellii* RNAi-mutant strains with WT (DPB079), Δ*ago1* (DPB325), Δ*dcr1* (DPB534), and Δ*xrn1* (DPB535) strains were performed by mixing equal volumes (100 μL) of saturated YPD cultures in a round-bottom 96-well plate and incubating at 30°C for 24 h. Diploid cells were then selected by plating these mixed saturated cultures on SD – Ade, – Lys agar plates (for RNAi mutants x DPB325) or SD – Met, – Lys agar plates (for all other matings). Flow cytometry was performed as with the haploid mutant strains, except with no gating for eGFP positive cells. Diploid strains presented in Figure 1D were created from matings of the WT genetic-selection strain (DPB537) and its derived RNAi-mutant strains with a WT strain (DPB079) or a Δ*xrn1* strain (DPB535) of the opposite mating type. The diploid *XRN1* knockout strain was created by mating strain DPB541 with strain DPB510.

### Flow cytometry of strains with RNA-decay mutations

Non-inducing SC (+ 2% glucose) 3 mL cultures were inoculated with strains that contained neither *GFP* nor a *GFP*-silencing construct (DPB211, DPB443, DPB442, DPB620, DPB601, DPB608), or contained *GFP* and an empty *GFP*-silencing construct (DPB331, DPB334, DPB337, DPB625, DPB469, DPB611), and 3 mL inducing SC (+ 2% galactose) cultures of strains that contained *GFP* and a *GFP*-silencing construct (DPB333, DPB336, DPB339, DPB629, DPB471, DPB613) and incubated at 25°C until they reached saturation. Strains with *DCP2* disrupted were inoculated before the other cultures, as they took > 24 h to reach saturation. Saturated cultures were diluted to OD_600_ 0.04 in 13 mL inducing or non-inducing media and grown until all cultures were in mid-log phase (OD_600_ 0.5 – 0.8). Flow cytometry was then performed on 500 μL of each culture as described for other strains. Strains used in Figure 1E were DPB211, DPB443, DPB442, DPB620, DPB331, DPB334, DPB337, DPB625, DPB333, DPB336, DPB339, DPB629, DPB601, DPB608, DPB469, DPB611, DPB471, DPB613.

### Serial-dilution spot assay

Inducing SC (+ 2% galactose) cultures of *N. castellii* strains were inoculated and incubated at 30°C for 12 h. These cultures were centrifuged for 5 min at 3000 x *g*, resuspended in SC (+ 2% galactose), and used to re-inoculate new cultures at OD_600_ 0.2. Cultures were grown until OD_600_ 1.0, centrifuged for 5 min at 3000 x g, and resuspended in water. Cells were diluted to 2 x 10^6^ colony forming units (CFU)/mL and subsequent tenfold dilutions were performed. Cells (5 μL) were plated on SC (+ 2% galactose) and SGal + Lys agar plates and grown for at least ten days. Strains from top to bottom in Figure 1C are DPB537, DPB541, DPB578, DPB582.

### Genome sequencing and analysis

Genomic DNA was isolated using as described (15) from saturated YPD cultures of *N. castellii* the genetic-selection strain (DPB537) and RNAi-mutant strains. DNA concentration was calculated using the Qubit dsDNA HS (High Sensitivity) Assay Kit (Invitrogen Q32854) with an Invitrogen Qubit Fluorometer. Genomic DNA libraries were prepared with either the Nextera DNA Library Preparation Kit (Illumina) or the QIAseq FX DNA Library Kit (QIAGEN 180475) and sequenced using Illumina SBS. Single-end sequencing was performed on the HiSeq platform, and paired-end sequencing was performed on the HiSeq or NextSeq platforms.

The *N. castellii* genome was downloaded from Yeast Genome Order Browser (YGOB - http://ygob.ucd.ie/). Sequences of the integrating plasmids containing *GFP*, *URA3* and *HIS3*-*PEST* along with their silencing constructs, as well as the integrating plasmids containing extra copies of *AGO1* and *DCR1* were inserted into the FASTA file. STAR (v.2.4) was used to create an indexed genome (16). Reads were aligned to the genome using STAR (v.2.4) with the parameters ‘--outFilterMismatchNmax 4 --clip3pAdapterSeq CTGTCTCTTATAC -- readMatesLengthsIn NotEqual --alignIntronMax 1 --alignMatesGapMax 1000’. Samtools (v1.3) ‘view’, ‘sort’, and ‘rmdup’ commands were used to sort the BAM file and remove duplicate reads (17). Samtools ‘mpileup’ command with the parameters ‘-t DP -t SP -ug’ was used to make a BCF file. Bcftools (v1.3) ‘call’ command with the parameters ‘-vm’ was used to call variants for each strain. A custom Python script was employed to remove the parental strain variants from each of the mutant strain variants, and to identify and select the variants that were called as homozygous by bcftools (as these were haploid strains). Custom Python scripts were used to characterize each of the remaining variants as being either genic (present in an ORF) or intergenic, and as being either exonic or intronic. Exonic ORF variants were further characterized as being either synonymous, nonsynonymous, or frameshift mutations, and the nonsynonymous mutations were classified as being either missense or nonsense mutations using custom Python scripts. The variants and the mutated genes in each of the mutant strains were compiled. Variants that occurred in more than two mutant strains were removed. All frameshift variants that occurred in repetitive tracts of DNA for which the variant caller (bcftools) identified more than one position of the repeat as being mutated were also removed. The remaining variants were considered bona fide mutations that occurred during either mutagenesis or selection of the mutant strains.

### Ago1p over-expression, immunoprecipitation and mass spectrometry

Overnight YPD cultures of diploid strains DPB201 and DPB202 were inoculated, grown to saturation, diluted to OD_600_ 0.05 in 250 mL YPD and grown until OD_600_ 1.0. Cultures were then induced by adding galactose to 2% and grown at 30°C for 20 h before harvesting to make 400– 500 mg pellets. An equal volume of lysis buffer (50 mM HEPES pH 7.6, 300 mM NaCl, 0.1 mM EDTA, 0.1 mM EGTA, 0.25% NP-40) was added to the pellets. Cells were lysed with acid-washed glass-bead beatings of 4 x 45 seconds with >4 min between bead beatings. 100 μL 50% anti-HA affinity matrix (Roche 11815016001) was added to the lysate and incubated 12 h at 4°C in experiment 1 and 60 μL of 50% anti-HA-agarose (Sigma A2095) was added and incubated for 2 h at 4°C in experiment 2. The affinity agarose was then washed four times with lysis buffer and all liquid was removed after the last wash with a needle. Proteins were eluted with 1x volume of HA peptide (1 μg/μL) with incubation at 37°C for 15 min in experiment 1 and with incubation at 4°C for 12 h in experiment 2. Liquid was removed from the beads, 2x SDS buffer was added and then boiled for 5 min and samples were run on a 4-12% Tris-Glycine gel.

A silver stain of the Tris-Glycine gel was performed using Pierce Silver Stain for Mass Spectrometry (ThermoFisher Scientific 24600). Each sample band in the gel was cut into ∼5 mm squares and was washed overnight in 50% methanol. These gel fragments were subsequently washed with a solution of 47.5% methanol, 5% acetic acid for 2 h and then dehydrated with acetonitrile and dried in a speed-vac. To reduce disulfide bonds, the dried gel pieces were incubated in 30 µL 100 mM ammonium bicarbonate with 10 mM dithiothreitol for 30 min and were then incubated in 30 µL of 100 mM ammonium bicarbonate and 100 mM iodoacetmide for cysteine alkylation. The gel fragments were then washed with acetonitrile, 100 mM ammonium bicarbonate and acetonitrile, in sequence, and dried in a speed-vac. The fragments were then rehydrated with a sufficient volume of 50 mM ammonium bicarbonate. To digest the proteins, trypsin was added (final concentration of 20 ng/µL), incubated on ice for 10 min, and then digested overnight at 37°C with gentle shaking. The digested peptides were extracted from the gel slices by sequential 10 min incubations at 37°C with shaking of the slices with 50 µL of 50 mM ammonium bicarbonate, then 50 µL of a solution of 47.5% acetonitrile, 5% water, and 5% formic acid, and finally 50 µL of acetonitrile/formic acid solution once again. The supernatants from each of these sequential incubations were pooled, and sample volumes were reduced to ∼15µL using a speed-vac concentrator.

Samples were analyzed by reversed-phase high-performance liquid chromatography (HPLC) using a Thermo EASY-nLC 1200 HPLC equipped with a self-packed Aeris 1.7 µm C18 analytical column (0.075 mm by 14 cm, Phenomenex). Peptides were eluted using standard reverse-phase gradients and analyzed using a Thermo Q Exactive HF-X Hybrid Quadrupole-Orbitrap mass spectrometer (nanospray configuration). The resulting fragmentation spectra were correlated against the known peptide database using Scaffold (Proteome Software Inc.). Analyses in Table S3 were performed using Scaffold.

### Ago1p-Xrn1p co-immunoprecipitation

Overnight YPD cultures of haploid strains DPB215, DPB220, DPB221 were grown, diluted to OD_600_ 0.2 in 150 mL YPD, and then grown until OD_600_ 0.8. Cultures were harvested by centrifugation, washed with 25 mL cold PBS, transferred to an Eppendorf tube in 750 μL cold PBS, and resuspended in 1 volume IP buffer (50 mM HEPES pH 7.6, 300 mM NaCl, 5 mM MgCl_2_, 0.1 mM EDTA, 0.1 mM EGTA, 5% glycerol, 0% NP-40, PMSF with 2 cOmplete protease inhibitor cocktail tablets (Roche 04693116001) per 50 mL of buffer). Cells were lysed with acid-washed glass bead beatings of 4 x 45 seconds with >4 min between bead beatings. The beads were removed via low speed centrifugation, 4 volumes of IP buffer were added, and cell debris was removed by centrifuging the extract at 21,000 x *g* for 5 sec. The extract was cleared by centrifuging twice at 10,000 x *g* for 5 min and retaining the supernatant. Absorbance readings (260 nm) of 1:10 dilutions of the lysate were performed with a Nanodrop (Thermo Scientific). The amount of extract needed for 30 A260 units for each lysate was aliquoted, IP buffer was added to 990 μL, and 10 μL RNase A/T1 cocktail (Ambion) or SUPERase-In was added. 0.3 A260 units of lysate was removed as the 1% input samples, the volume to was raised to 10 μL with IP buffer, and then incubated at 4°C for 12 h. 20 μL of EZview Red ANTI-FLAG M2 Affinity Gel (Sigma F2426) was added to lysates from strain DPB215 +/– RNase and DPB221 or 20 μL of EZview Red Anti-HA Affinity Gel (Sigma E6779) was added to lysates from strain DPB215 +/– RNase and DPB220. The lysates were incubated at 4°C for 12 h. The affinity agarose was sedimented by centrifugation, and the supernatant was removed. For the analysis to confirm RNA degradation, 3 A260 = 100 μL (10%) of supernatant was saved, and additionally 10 μL of the supernatant was saved for 1% supernatant samples. The affinity agarose was washed four times with IP buffer (each wash, 800 μL centrifuged 10,000 x *g* for 5 min), and the residual buffer was removed with a needle. 100 μL 1x Laemmli buffer (Bio-Rad 1610737) + 2.5% beta-mercaptoethanol (bME) was added to the affinity agarose for each sample. 10 μL 2x Laemmli buffer + 2.5% bME and 80 μL 1x Laemmli buffer + 2.5% bME was added to the input and supernatant samples. All samples were incubated at 90°C for 5 min, and then were centrifuged at room temperature for five min at 10,000 x *g*. All samples were run on a 4–12% Bis-Tris gel in MOPS buffer. The proteins were transferred from the gel to a PVDF membrane (Invitrogen LC2002), each blot was probed with anti-HA7 (1:2000) (Sigma H3663) and anti-Flag BioM2 (1:2000) (Sigma F9291) overnight at 4°C, and probed with the secondary antibody anti-mouse IgG HRP (1:5000) (Amersham NXA391) for 1–2 h at room temperature. The blots were stripped with 100 mM bME, 2% SDS, 62.5 mM Tris pH 6.8 at 50°C for 30 min, shaking every 10 min. For the RNA-degradation analysis, samples containing 100 µl of supernatant were phenol extracted, precipitated, and resuspended in 45 μL water. 1 μL of the RNA sample was added to 2 μL water, and then 3 μL 2x glyoxal loading dye was added. Samples were then incubated at 50°C for 30 min to denature the RNA and then placed on ice. Samples were resolved on a 1% agarose gel run in NorthernMax-Gly Gel Prep/Running Buffer (Invitrogen AM8678) and visualized with a ChemiDoc MP Visualization System (Biorad).

### Ago1p, Dcp2p, Xrn1p in vivo imaging

YPD cultures of strains DPB231 and DPB234 were grown overnight at 25°C. These overnight cultures were split in two, harvested by centrifugation, washed with CSM with or without glucose, resuspended in CSM with or without glucose, and grown at 25°C for 1 h before imaging. 488-nm and 543-nm lasers on a Zeiss LSM 510 confocal microscope were used to excite GFP and mCherry, respectively. Images were edited with ImageJ (18).

### Small-RNA blots

Total RNA was isolated from log-phase cultures (OD_600_ 0.8-1.0) of *N. castellii* using the hot-phenol method. 20 μg of total RNA was loaded per lane of a denaturing 15% polyacrylamide gel. After blotting, carbodiimide was used to crosslink small RNAs to the membrane (19). DNA probes (*N. castellii* palindrome-derived siRNA, 5′-CTATCTTCATCGATTACCATCTA; *N. castellii* Y′ siRNA, 5′-TCATGGTTAAGTATGGACGTCAA; *N. castellii* U6 small nuclear RNA, 5′-TATGCAGGGGAACTGCTGAT; *H. sapiens* miR-21, 5′-TCAACATCAGTCTGATAAGCTA) were labeled at their 5′ termini, as was the LNA probe for *A. thaliana* miR156 (20). Strains used in Figure 3C were DPB220 and DPB622. Strains used in Supplemental Figure 1A were DPB211, DPB443, DPB442, DPB620, DPB601, DPB608. Periodate oxidation and beta-elimination was as described (21).

### Small-RNA sequencing, RNA-seq, and analysis

#### Sequencing libraries

Total RNA was isolated from *N. castellii* strains DPB220, DPB228, DPB622, DPB537, and DPB541. For small-RNA sequencing, 0.5 nM of four small RNA oligonucleotides (xtr miR-427, 5′-GAAAGUGCUUUCUGUUUUGGGCG; dme miR-14, 5′-GGGAGCGAGACGGGGACTCACT; Synthetic_siRNA_1_Guide, 5′-UAGUGCAGGUAGGUAUUUUUGUU; Synthetic_siRNA_1_Passenger, 5′-CAAAAAUACCUACCUGCACUAUA) were added to 10 μg of total RNA for use as internal standards. Small-RNA cDNA libraries were prepared as described (22), except 9–26-nt RNAs were isolated and sequenced, using 9-nt and 26-nt RNA radiolabeled standards (5′-AAACCAGUC and 5′ -AGCGUGUACUCCGAAGAGGAUCCAAA) to follow ligations and purifications.

For RNA-seq, 0.5 ng of two mRNAs (chloramphenicol and firefly luciferase) were added to 5 μg of total RNA for use as internal standards. Ribosomal RNA was depleted from the samples using Ribo-Zero Gold rRNA Removal Kit (Yeast) (Illumina, MRZY1324). RNA cDNA libraries were prepared using NEXTflex rapid directional mRNA-seq kit (Bioo Scientific, NOVA-5138-10). Libraries were sequenced using the Illumina SBS platform.

#### Genome and small-RNA cluster assembly

The *N. castellii* genome used for analysis of small-RNA and RNA sequencing from wild-type (DPB220), *AGO1* slicing-impaired (DPB228), and Δ*xrn1* (DPB622) strains in Figure 3 was downloaded from YGOB. The sequences of the Y′-consensus ORF (1) as well as the chloramphenicol and firefly luciferase internal-standard mRNA sequences were added to the FASTA file. The *N. castellii* genome used for analysis of RNA sequencing from wild-type (DPB537) and Δ*xrn1* (DPB541) genetic-selection strains in Supplemental Figures 1 and 4C was the *N. castellii* selection genome used for genome-sequencing analysis supplemented with the Y′-consensus ORF as well as the chloramphenicol and luciferase internal-standard mRNA sequences. The *N. castellii* genome used for RNA sequencing analysis with strains DPB537 and DPB541 in Supplemental Figure 4D was downloaded from Ensembl (ASM23734v1). STAR (v.2.4) was used to generate indexed genomes.

The coordinates of annotated small-RNA clusters (1) were converted from scaffolds of the unfinished *N. castellii* genome to the genomic coordinates of the finished *N. castellii* genome using the liftOver command line tool from UCSC (https://genome.ucsc.edu/). The sequence of one annotated small-RNA cluster did not match the finished genome, and thus this cluster was omitted. For some small-RNA clusters, the former scaffold coordinates overlapped in the finished genome; these were combined to generate larger clusters in the updated set of small-RNA clusters. In instances in which a region of a previously annotated small-RNA cluster was partially deleted in the new genome, the remainder was retained as a smaller cluster. In cases in which the scaffold sequence of a small-RNA cluster mapped to multiple sequences in the finished genome, only one set of coordinates for the cluster was retained. Several new clusters were also identified as genomic regions for which 22–23-nt reads mapped but were not previously annotated as small-RNA clusters and did not overlap known rRNA or tRNA loci. The updated table of the genomic coordinates of the small-RNA clusters is provided (Table S4).

#### Small-RNA read processing and expression analysis

The first four nucleotides of each sequencing read were removed using fastx_trimmer from the FASTX-toolkit (http://hannonlab.cshl.edu/fastx_toolkit/index.html). The 3′ adapter was then trimmed using cutadapt (v1.8) (23) with parameters ‘-a NNNNTCGTATGCCGTCTTCTGCTTG -m 9’. High-quality reads were then selected using fastq_quality_filter from the FASTX-toolkit with parameters ‘-v -q 30 -p 100 -Q 64 -z’. 9–26-nt, and 10–12-nt or 22–23-nt high-quality reads were selected using cutadapt. Reads were aligned to their appropriate *N. castellii* genomes using STAR (v.2.4) allowing no mismatches or gaps with the parameters ‘--runThreadN 30 --alignIntronMax 1 --alignIntronMin 2 --scoreDelOpen - 10000 --scoreInsOpen -10000 --outFilterMismatchNmax 0 --seedSplitMin 9 --outFilterType BySJout --outFilterMultimapNmax 1000 --outSAMtype BAM SortedByCoordinate -- readFilesCommand zcat --outFilterIntronMotifs RemoveNoncanonicalUnannotated -- alignEndsType EndToEnd’. To remove rRNA and tRNA reads, featureCounts (24) was used to fractionally count the number of times each read mapped to annotated rRNA loci and their intergenic regions (Ncas_Chr3: 572204–588441) and to annotated tRNA loci in the *N. castellii* genome using the parameters ‘-s 1 -f -O -M --fraction --donotsort’. Additionally, featureCounts was used to produce a SAM file of these rRNA and tRNA reads which were then removed from the original files of 9–26-nt, 10–12-nt or 22–23-nt high quality reads using custom Python scripts. The high-quality reads that were depleted of rRNA and tRNA reads were then aligned to the appropriate *N. castellii* genomes using STAR (v.2.4) with the same parameters as before. To fractionally count the number of times each read mapped to specific features in the *N. castellii* genome, featureCounts (24) was utilized using the parameters ‘-s 1 -f -O -M --fraction -- donotsort’. For the small-RNA sequencing analysis with wild-type (DPB220), *AGO1* slicing-impaired (DPB228), and Δ*xrn1* (DPB622) strains, this fractional counting was performed using either a GTF file comprised of the entire genome divided into genic or intergenic regions or a GTF file of the previously annotated small-RNA clusters, each of which included the Y′-consensus ORF. For the small-RNA sequencing analysis with wild-type (DPB537) and Δ*xrn1* (DPB541) genetic-selection strains, this fractional counting was performed using a GTF file of the previously annotated small-RNA clusters, including the Y′-consensus ORF as well as the synthetic hairpins used in the genetic selection. Fractional counts were converted to reads per million for each locus and then normalized based on the internal small-RNA standard reads in each library. In Figure 3B, the 22–23-nt reads that mapped to the small-RNA clusters were extracted from the BAM file and categorized based on whether they mapped exclusively to palindromic or non-palindromic loci or if they mapped to both types of loci.

#### Analysis of passenger-strand stability

Analysis started with 22–23-nt and 10–12-nt reads that mapped to the small-RNA clusters. Custom Python scripts identified 22–23-nt reads that could pair perfectly with each other with the 2-nt 3′ overhangs characteristic of siRNA duplexes, as well as 10- or 11-nt reads that perfectly matched 5′ ends of these species and 12-nt reads that perfectly matched the 3′ ends of these species. Ambiguous pairs for which a 22–23-mer had multiple 22–23-mers that could potentially pair to it but with different 2-nt 3′ overhangs were excluded. In instances for which a 10–12-mer matched more than one 22–23-mer, its number of reads was divided by the number of matching 22–23-mers and assigned equally to these 22–23-mers. Each guide-passenger pair displayed in Figure 5 passed the following cutoffs in the wild-type strain, the Δxrn1 strain, as well as the AGO1 slicing-impaired strain: the guide strand had ≥100 reads; the full-length passenger strand had ≥1 read; and at least one of the two passenger-strand cleavage fragments had ≥1 read. Additionally, the ratio of full-length passenger-strand reads to guide-strand reads was greater in the *AGO1* slicing-impaired strain than in the WT strain, and the ratio of cleaved passenger-strand reads to guide-strand reads was greater in the WT strain than in the *AGO1* slicing-impaired strain.

#### RNA-seq read processing and expression analysis

RNA-seq reads were aligned to the genome using STAR (v.2.4) with the parameters ‘-- outFilterType BySJout --outFilterMultimapNmax 1000 --outFilterMismatchNoverReadLmax 0.04 --alignIntronMax 1100 --outSAMtype BAM SortedByCoordinate --readFilesCommand zcat --outFilterIntronMotifs RemoveNoncanonicalUnannotated’. As with the small-RNA sequencing, featureCounts were used to fractionally count the number of times each read mapped to the genome using the parameters ‘-s 2 -f -O -M --fraction --donotsort’. This fractional counting was performed using a GTF file of the previously annotated small-RNA clusters, which included the Y′-consensus ORF. The fractional counts were converted to transcripts per million and normalized in the same manner as the small-RNA sequencing libraries. Browser tracks for visualizing RNA-seq results shown in Supplemental Figure 4C were made using IGV (v.2.4.10) (25,26). The metagene profile shown in Supplemental Figure 4D was generated using ngs.plot.r (v.2.61) (27) using parameters ‘-R genebody -SE 0 -L 100’.

#### RNAs for in vitro assays

The guide RNA (5′-UAAAGUGCUUAUAGUGCAGGUAG), the passenger-strand RNA (5′-ACCUGCACUAUAAGCACUUUAAG), the passenger-strand RNA for 3′ labeling (5′-ACCUGCACUAUAAGCACUUUAA), and the slicing target (5′-GGGAGAAACAAAAAUACCUACCUGCACUAUAAGCACUUUACCAUCUCAAACUUACUCAGA) were synthesized, purified, and labeled as described (28) with the following adaptations. Capping and cap-labeling were performed with the Vaccinia Capping System (NEB). To generate unlabeled capped target, a large-scale capping reaction (60 μL reaction containing 60 μg target RNA) was performed and capped RNA was gel purified on a long, denaturing 12% polyacrylamide gel that resolved the capped and uncapped species.

#### Ago1p and Xrn1p protein purification

50 mL SR – Ura cultures (S – Ura media + 2% raffinose) were inoculated with cells from SD – Ura plates of *S. cerevisiae* strains DPB598 (which expressed WT *N. castellii* Ago1p in a Δ*xrn1* background), DPB1100 (which expressed WT *N. castellii* Xrn1p in a Δ*xrn1* background), DPB1101 (which expressed mutant *N. castellii* Xrn1p, containing D206N and D208N substitutions at the active site in a Δ*xrn1* background). 1000 mL liquid yeast extract, peptone (YEP) + 2% raffinose cultures were inoculated with cells from the 50 mL SR – Ura cultures at OD_600_ ∼0.1. After growth at 30°C to OD_600_ 1.2, 100 mL of 20% galactose was added, and the cultures were incubated with shaking at 30°C for 24 h. Cultures were harvested by centrifugation at 2000 x *g* for 5 min. The cells were washed once with 40 mL 0.1 mM 4-(2-aminoethyl)benzenesulfonyl fluoride hydrochloride (AEBSF, Sigma-Aldrich A8456), transferred into conical 50 mL tubes and centrifuged at 1700 x *g* for 5 min. The cells were washed once in 300 mM glutamate/sorbitol buffer (50 mM Hepes pH 7.6, 1 mM EGTA, 1 mM EDTA, 5 mM magnesium acetate, 10% glycerol, 300 mM potassium glutamate, 800 mM sorbitol, 1 mM DTT added fresh) and pelleted by centrifugation at 1500–1700 x g for 5–7 min, and then 1/3 cell volume of glutamate buffer with protease inhibitor (50 mM Hepes pH 7.6, 1 mM EGTA, 1 mM EDTA, 5 mM magnesium acetate, 10% glycerol, 300 mM potassium glutamate, 1 mM DTT added fresh, 1 cOmplete, Mini, EDTA-free Protease Inhibitor Cocktail (Roche 11836170001) tab per 10 mL buffer) was added. The cell slurry was dripped slowly into liquid nitrogen to form frozen beads of cells, which were stored at –80°C. The cells were lysed in a SPEX

SamplePrep 6870 Freezer/Mill (10 cycles of 2 min grinding and 2 min cool down). After cell lysate was thawed on ice for 60 min, glutamate buffer with protease inhibitor was added (∼5 mL for 5 g of cell powder) and lysate was centrifuged at 30,000 x g for 90 min at 4°C. The supernatant was collected and centrifuged at 30,000 x g for 30 min and then additional glutamate buffer with protease inhibitor was added (another 5 mL for 5 g of cell powder). The supernatant was added to ANTI-FLAG M2 Affinity Gel (Millipore A2220; adding 2 mL gel slurry for 5 g of cell powder) and incubated with rotation for 2.5 h at 4°C. The affinity agarose was washed twice with 1M glutamate/NP40 buffer (50 mM Hepes pH 7.6, 1 mM EGTA, 1 mM EDTA, 5 mM magnesium acetate, 10% glycerol, 1 M potassium glutamate, 0.01% NP-40, 5 mM DTT added fresh) and twice with 300 mM glutamate/NP40 buffer (50 mM Hepes pH 7.6, 1 mM EGTA, 1 mM EDTA, 5 mM magnesium acetate, 10% glycerol, 300 mM potassium glutamate, 0.01% NP-40, 5 mM DTT added fresh). Proteins were eluted by incubation of the affinity agarose with 300 mM glutamate buffer containing TEV protease (50 mM Hepes pH 7.6, 1 mM EGTA, 1 mM EDTA, 5 mM magnesium acetate, 10% glycerol, 300 mM potassium glutamate, 0.01% NP-40, 5 mM DTT added fresh, 25 μL TEV protease (Sigma-Aldrich T4455) per 1 mL buffer) for 15 h at 4°C. For the Ago1p purification, the TEV protease was removed using Ni-NTA Agarose (QIAgen, 30210) according to manufacturer’s protocol. Concentrations of the purified proteins were determined by Bradford Assay using the Bio-Rad Protein Assay Dye Reagent Concentrate (Bio-Rad 5000006) and calculating the ratio of the samples’ absorbance at 595 nm and 472 nm. Protein aliquots were flash frozen in liquid nitrogen and stored at –80°C.

### RISC purification

Lysate of *S. cerevisiae* strain DPB598, prepared as described above, was incubated with an siRNA duplex containing 5′-radiolabeled miR-20a and the corresponding passenger strand (28) (final concentration, 0.25 μM) for 7 h at 30°C with shaking (600 RPM). Subsequent purification of miR-20a-loaded Ago1p-RISC was based on affinity to the miR-20a seed (29). Streptavidin agarose beads (Sigma-Aldrich S1638) were resuspended in storage buffer and 1.5 mL of slurry was washed three times with 300 mM glutamate/NP40 buffer and bound to 5 nmol capture 2′*-O-* methyl RNA oligonucleotide (5′-CUCACCUUCUACACCACC**GCACUUUA**UCCUUACACAC-3′-biotin; bold, match to the miR-20a extended seed region) in 1.5 mL of water by incubating for 90 min at 30°C. The agarose beads were then washed three times with 300 mM glutamate/NP40 buffer, incubated with lysate for 10 h at 4°C, and then washed three times with 1M glutamate/NP40 buffer and three times with 300 mM glutamate/NP40 buffer. Ago1p-RISC was then eluted by adding 10 nmol competitor DNA oligonucleotide (5′-AAGGATAAAGTGCGGTGGTGTAGAAGGTGAG-3′) in 1 mL 300 mM glutamate/NP40 buffer to the agarose beads. After incubating for 2 h at 4°C, eluate was collected, and Ago1p-RISC was bound to and eluted from ANTI-FLAG M2 Affinity Gel as described above. Ago1p-RISC concentration was determined based on the fraction of the input radioactivity that remained associated with purified Ago1p-RISC. Protein aliquots were flash frozen in liquid nitrogen and stored at –80°C.

### siRNA-binding assay

Proteins and RNAs were diluted in siRNA-binding buffer (50 mM Hepes pH 7.6, 1 mM EGTA, 1 mM EDTA, 5 mM magnesium acetate, 10% glycerol, 300 mM potassium glutamate, 10 mM NaCl, 0.01% NP-40, 5 mM DTT added fresh). 10X RNA mix (10 nM miR-20a siRNA duplex with a 5′-radiolabeled guide strand, 100 nM unlabeled miR-20a siRNA duplex, 1 μM capped target RNA, 1 µM tRNA from baker’s yeast (Sigma-Aldrich R5636), 1 μL SUPERase-In RNase Inhibitor (Invitrogen AM2696) per 20 μL RNA mix) was made, and 1.1 μL of this mix was added to a 1.5 mL G-tube siliconized microcentrifuge tube (BIO PLAS Inc., 4165SL). 1.1X protein mixes contained either no protein, 222 nM mut. Xrn1p, 222 nM WT Xrn1p, 111 nM WT Ago1p, 111 nM WT Ago1p and 222 nM mut. Xrn1p, or 111 nM WT Ago1p and 222 nM WT Xrn1p. Each protein mix was pre-incubated at 30°C for 5 min before adding 9.9 µL of the protein mix to the RNA mix. After incubation at 30°C for the indicated time, 10 μL of each reaction was removed and analyzed using layered nitrocellulose–nylon filter-binding assay (30). Radiolabelled RNA was visualized by phosphorimaging (Fujifilm BAS-2500) and quantified using Multi Gauge (Fujifilm) software.

### Passenger-strand-cleavage and target-slicing assays

For both assays, proteins and RNAs were diluted in the siRNA-binding buffer. For the combined passenger-strand and target-slicing assay in Figure 4A–D, 5X RNA mix (5 nM miR-20a duplex with a 5′-radiolabeled passenger strand, 50 nM unlabeled miR-20a siRNA duplex; ∼500 nM cap-radiolabeled target RNA, 500 nM capped target RNA, 500 nM tRNA from baker’s yeast, and 1 μL SUPERase-In RNase Inhibitor per 20 μL RNA mix) was made, and 2 μL of this RNA mix was added to a 1.5 mL G-tube siliconized microcentrifuge tube. 1.25X protein mixes contained either no protein, 250 nM mut. Xrn1p, 250 nM WT Xrn1p, 125 nM WT Ago1p, 125 nM WT Ago1p and 250 nM mut. Xrn1p, or 125 nM WT Ago1p and 250 nM WT Xrn1p. Each protein mix was pre-incubated at 30°C for 5 min before adding 8 µL of the protein mix to the RNA mix. After incubation at 30°C for the indicated time, 2 μL of the reaction was removed and quenched by adding to 5 μL of urea loading buffer (8M urea pH 8.0, 25 mM EDTA, 0.025% xylene cyanol, 0.025% bromophenol blue) and 3 μL water. For the assay of passenger-strand cleavage in Supplemental Figure 2A–C, the reactions were carried out as above, except no target was added to the RNA mix and the miR-20a siRNA duplex contained a 3′-cordycepin-labeled passenger strand.

For the Ago1p-RISC target-slicing assay in Figure 6A–C, 10X RNA mix (10 nM cap-radiolabeled target RNA, 1 µM capped target RNA, 1 µM tRNA from baker’s yeast, and 1 μL SUPERase-In RNase Inhibitor per 20 μL RNA mix) was made, and 1 μL of this RNA mix was added to a 1.5 mL G-tube siliconized microcentrifuge tube. 1.1X protein mixes contained either no protein, 11.1 nM Ago1p-RISC, 11.1 nM Ago1p-RISC and 11.1 nM mut. Xrn1p, 11.1 nM Ago1p-RISC and 11.1 nM WT Xrn1p, 11.1 nM Ago1p-RISC and 33.3 nM mut. Xrn1p, 11.1 nM Ago1p-RISC and 33.3 nM WT Xrn1p, 11.1 nM Ago1p-RISC and 111.1 nM mut. Xrn1p, or 11.1 nM Ago1p-RISC and 111.1 nM WT Xrn1p. Each of the protein mixes was pre-incubated at 30°C for 20 min, before adding 9 µL of the protein mix to the RNA mix. After incubation at 30°C for the indicated time, 2 μL of each reaction was removed and quenched as for the combined assay. For the Ago1-RISC target-slicing assay in Figure 6D–F, the reactions were carried out as above, except the target was 3′-cordycepin-labeled.

To assess cleavage and slicing, RNAs were resolved on denaturing (7.5 M urea) 20% polyacrylamide gels, and radiolabeled cleavage products were visualized and quantified by phosphorimaging as above. At each time point, the fraction product was measured as F_p_ = product / (product + substrate). For Figure 6F, the fraction product was measured as F_p_ = product / (product + substrate + degradation intermediate). For Figure 6C, the multiple-turnover slicing data were fit in Prism 8 (GraphPad) to the burst and steady-state equation (31,32)

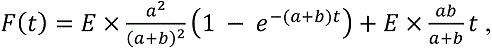

Where *F(t)* is target cleaved over time, *E* is the enzyme concentration, and *a* and *b* are rate constants (reported as *k*_1_ and *k*_2_, respectively) according to the following scheme,

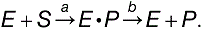

## Supplemental Figure Legends

**Supplemental Figure 1.**
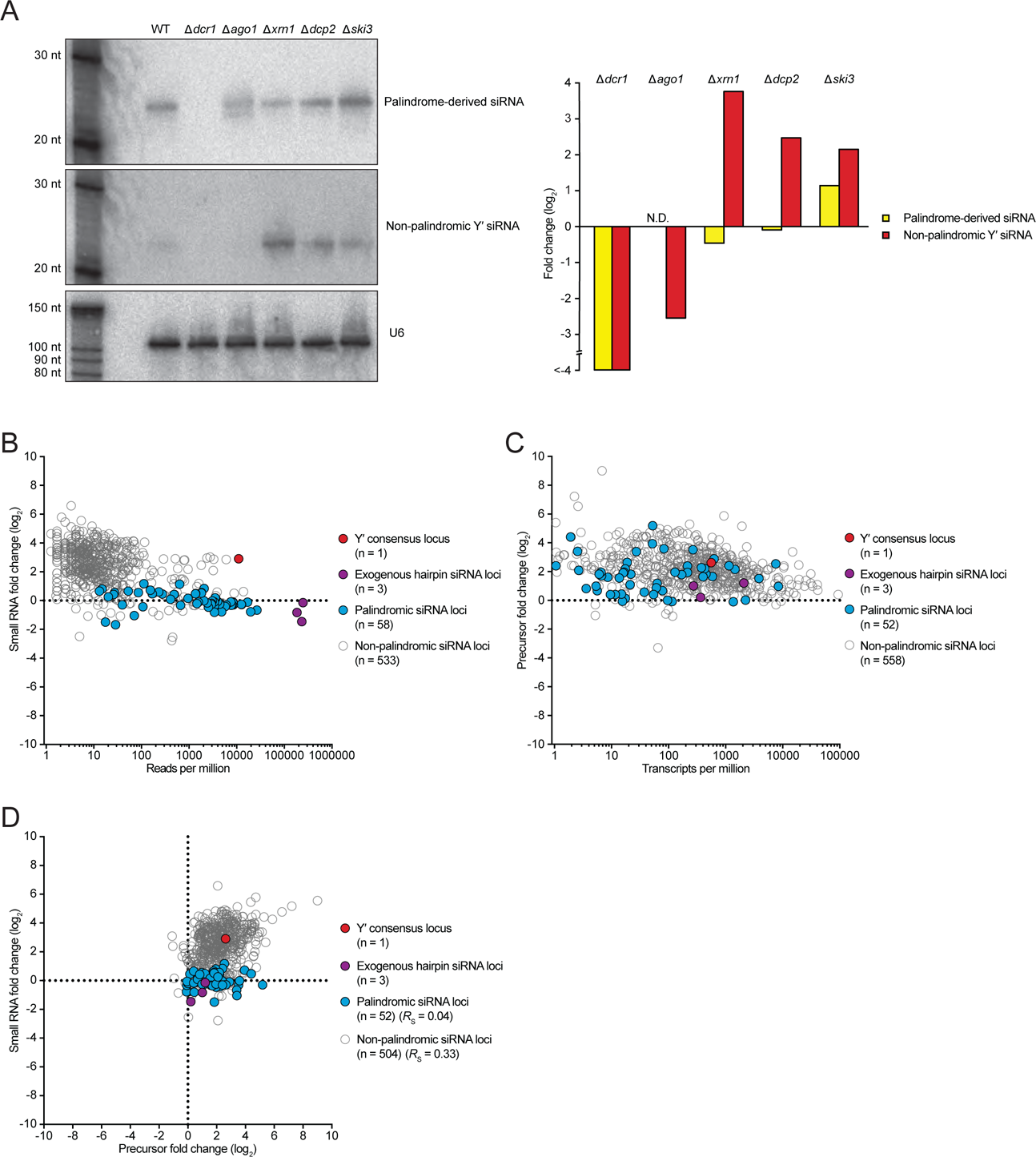
The effects of Xrn1p and other proteins on the abundance of small RNAs and their precursors in *N. castellii*. **A)** Effects of RNAi proteins and several RNA-decay factors on the expression levels of both a palindrome-derived siRNA and a non-palindrome-derived siRNA. Shown are the results of successively probing an RNA blot, as in **Figure 3C**. **B)** Effect of Xrn1p on the abundance of small RNAs originating from siRNA-producing loci of the *N. castellii* genetic-selection strain. Otherwise, as in **Figure 3A**. **C)** Effect of Xrn1p on the abundance of small-RNA precursor transcripts in the *N. castellii* genetic-selection strain. Otherwise, as in **Figure 3D**. **D)** Relationship between the changes in small RNAs and the changes in siRNA precursor transcripts observed after deleting *XRN1* in the *N. castellii* genetic selection strains. Otherwise, as in **Figure 3E**.

**Supplemental Figure 2.**
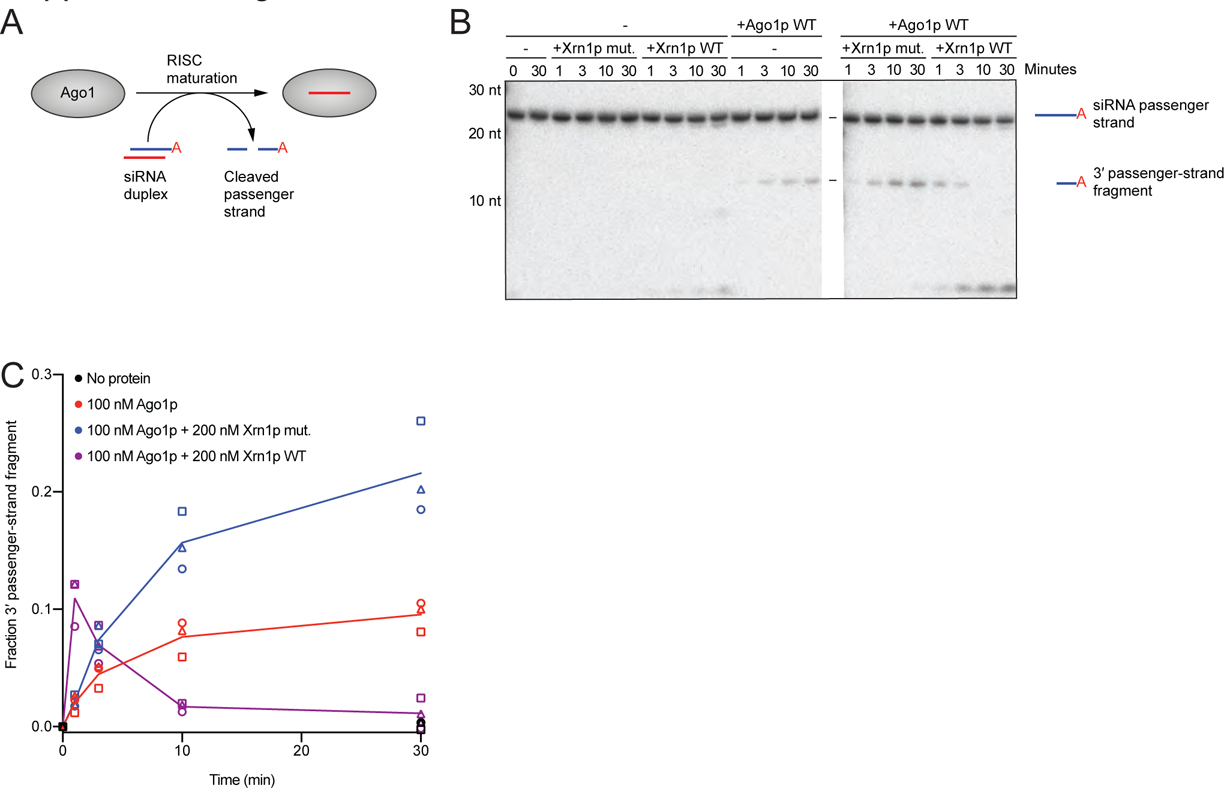
Impact of Xrn1p on stability of the 3′ passenger-strand cleavage fragment. **A)** Experimental scheme of the assay of duplex loading and passenger-strand cleavage. The red A in the duplex indicates a radiolabeled cordycepin at the 3′-end of the passenger strand. The initial concentration of siRNA duplex was 10 nM. **B)** Effect of Xrn1p on passenger-strand cleavage and degradation in the assay schematized in **A**. Shown is a representative denaturing gel resolving 3′-labeled passenger strand from its cleavage product after incubation with or without purified Ago1p (100 nM) and with or without Xrn1p (WT or mut.) (200 nM) for the indicated amount of time. **C)** Quantification of the 3′ passenger-strand fragment. Otherwise, as in **Figure 4C**.

**Supplemental Figure 3.**
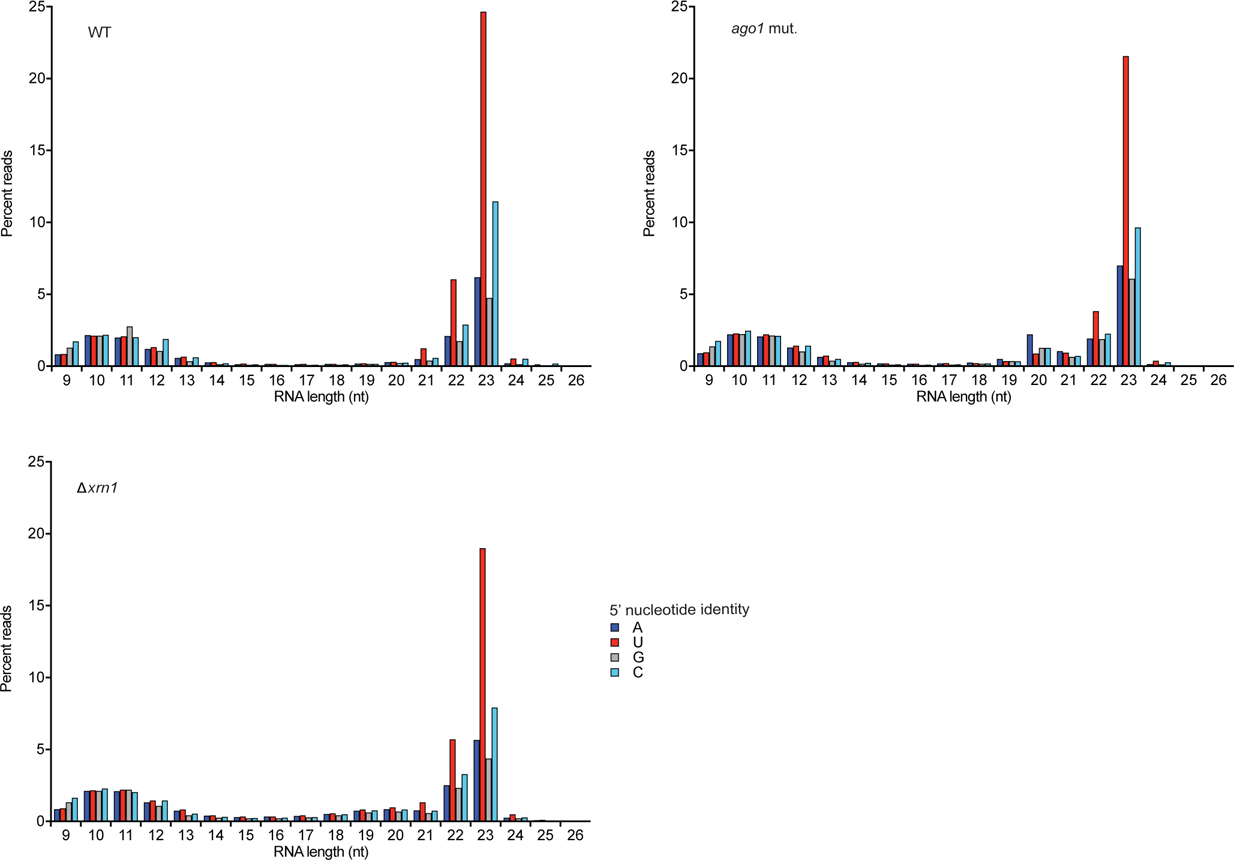
The length distribution of small-RNA reads from WT, slicing-impaired *ago1* mutant, and Δ*xrn1 N. castellii* strains. At each length from 9–26 nt, the number of genome-matching sequencing reads with the indicated 5′ nucleotide is plotted. Reads mapping to either rRNA or tRNA loci were excluded.

**Supplemental Figure 4.**
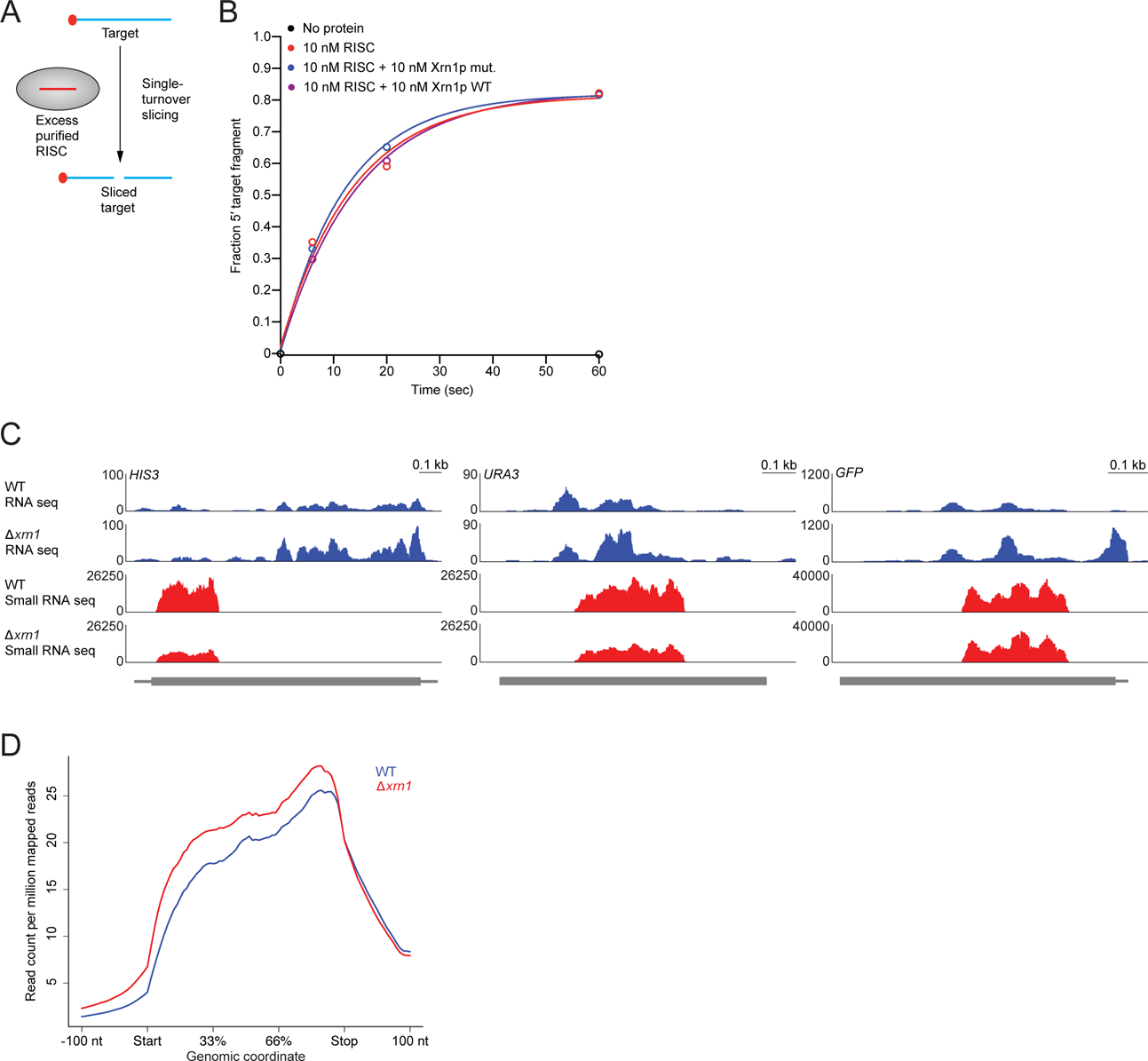
Effect of Xrn1p on single-turnover slicing and on accumulation of 3′ fragments of RNAi targets in vivo. **A)** Experimental scheme of single-turnover slicing of a cap-labeled target. The initial target concentration is substantially below that of RISC, otherwise as in **Figure 6A**. **B)** Quantification of the 5′ product of slicing under single-turnover conditions, as schematized in **A**. Slicing was carried out in single-turnover conditions, incubating cap-labeled target (1 nM) with purified RISC–miR-20a (10 nM), with or without Xrn1p (WT or mut.) (10 nM) for the indicated amount of time. Results are shown for one experiment with either no protein (black), RISC–miR-20a only (red), RISC–miR-20a with Xrn1p mut. (blue), and RISC–miR-20a with Xrn1p WT (purple). Fraction cleaved was calculated by dividing the signal of product / (product + substrate). Lines represent the best fit to the equation for one-phase association. **C)** Effects of Xrn1p on mRNAs targeted by RNAi in the genetic selection and on siRNAs targeting these mRNAs. Strand-specific RNA-seq (blue) and small-RNA sequencing (red) profiles are shown for mRNAs targeted by RNAi (*GFP*, *HIS3, URA3*) in WT and Δ*xrn1* genetic-selection strains. The mRNAs are schematized below the profiles, with thick gray lines indicating coding sequence and thin gray lines indicating untranslated regions. Track heights were normalized based on reads that mapped to the internal standards, with the RNA-seq reads for the WT strain normalized with respect to reads for the Δ*xrn1* strain and the small-RNA sequencing reads for the Δ*xrn1* strain normalized with respect to those of the WT strain. **D)** The effect of Xrn1p on the mRNAs of non-RNAi targets. Shown are metaplots of RNA-seq reads mapping to mRNAs that were not targets of RNAi in WT (blue) and Δ*xrn1* (red) genetic-selection strains. The start and stop codons are indicated, as are positions 100 nucleotides (nt) upstream and downstream of the ORF. Reads displayed are normalized to library size.

**Supplemental Figure 5.**
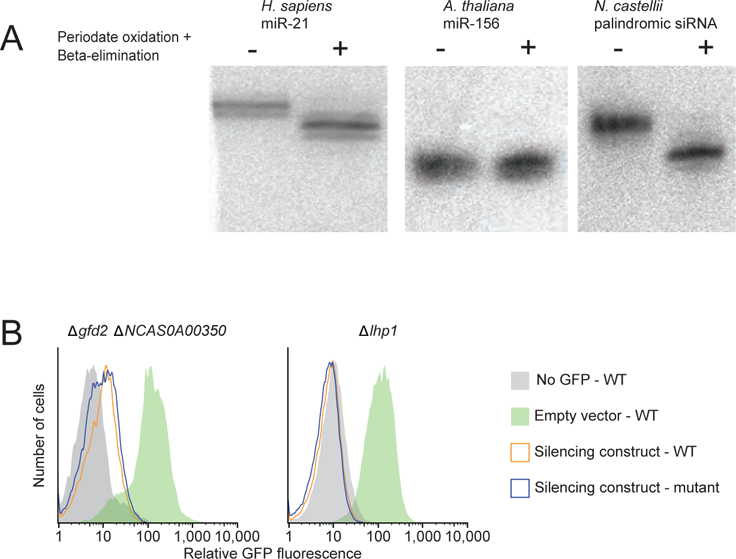
Assessment of siRNA 2′-O-methylation and the effect of disrupting RNAi cofactor orthologs in *N. castelii*. **A)** Examination of 2′-O-methylation of the 3′-terminal nucleotide of *N. castellii* siRNAs. Total RNA from a human cell line (HeLa), *Arabidopsis thaliana* leaves, and *N. castellii* was subjected to periodate oxidation and beta-elimination. Samples with and without treatment were then analyzed on RNA blots. The human sample was probed for miR-21, which is not 2′-O-methylated and is therefore susceptible to oxidation and elimination, the *A. thaliana* was probed for miR-156, which is 2′-O-methylated and therefore protected from oxidation and elimination, and the *N. castellii* sample was probed for a palindromic siRNA (as in **Figure 3C**). **B)** Effect of disrupting the orthologs of genes reported to enhance RNAi in other species. Shown are histograms of GFP fluorescence measured by flow cytometry in Δ*gfd2* Δ*NCAS0A00350* and Δ*lhp1 N. castellii* strains with the indicated GFP-silencing constructs. All strains were induced.

**Table S1.** Protein-coding or tRNA mutations identified in each mutant strain sequenced. For each mutation, the gene, the mutant sequence, the genomic coordinate, and the reference sequence are listed.

**Table S2.** Genes mutated in the mutant strains that were sequenced. For each gene, the number of sequenced strains that had mutations in the gene is indicated, as well as the total number of mutations and the number of unique and nonsense mutations.

**Table S3.** Ago1p immunoprecipitation and mass spectrometry results for two separate experiments. The tables were each sorted by enrichment of total spectra in the pull-down condition over the control condition.

**Table S4.** Updated genomic coordinates of siRNA-producing transcripts.

**Table S5.** Strains used and generated in this study.

**Table S6.** Plasmids generated in this study.

